# Thermogenic hydrocarbons fuel a redox stratified subseafloor microbiome in deep sea cold seep sediments

**DOI:** 10.1101/2020.02.02.928283

**Authors:** Xiyang Dong, Jayne E. Rattray, D. Calvin Campbell, Jamie Webb, Anirban Chakraborty, Oyeboade Adebayo, Stuart Matthews, Carmen Li, Martin Fowler, Adam Macdonald, Ryan A. Groves, Ian A. Lewis, Scott H. Wang, Daisuke Mayumi, Chris Greening, Casey R.J. Hubert

## Abstract

At marine cold seeps, gaseous and liquid hydrocarbons migrate from deep subsurface origins to the sediment-water interface. Cold seep sediments are known to host taxonomically diverse microorganisms, but little is known about their metabolic potential and depth distribution in relation to hydrocarbon and electron acceptor availability. In this work, we combined geochemical, metagenomic and metabolomic measurements in distinct sediment redox regimes to profile microbial activities within the uppermost 350 cm of a newly discovered cold seep in the NW Atlantic deep sea (2.3 km water depth). Depth-resolved metagenomic profiling revealed compositional and functional differentiation between near-surface sediments (dominated by Proteobacteria) and deeper subsurface layers (dominated by Atribacteria, Chloroflexi, Euryarchaeota and Lokiarchaeota). Metabolic capabilities of community members were inferred from 376 metagenome-assembled genomes spanning 46 phyla (including five novel candidate phyla). In deeper sulfate-reducing and methanogenic sediments, various community members are capable of anaerobically oxidizing short-chain alkanes (alkyl-CoM reductase pathway), longer-chain alkanes (fumarate addition pathway), and aromatic hydrocarbons (fumarate addition and subsequent benzoyl-CoA pathways). Geochemical profiling demonstrated that hydrocarbon substrates are abundant in this location, thermogenic in origin, and subject to biodegradation. The detection of alkyl-/arylalkylsuccinate metabolites, together with carbon isotopic signatures of ethane, propane and carbon dioxide, support that microorganisms are actively degrading hydrocarbons in these sediments. Hydrocarbon oxidation pathways operate alongside other deep seabed metabolisms such as sulfide oxidation, hydrogen oxidation, carbon fixation, fermentation and reductive dehalogenation. Upward migrated thermogenic hydrocarbons thus sustain diverse microbial communities with activities that affect subseafloor biogeochemical processes across the redox spectrum in deep sea cold seeps.

## Introduction

Marine cold seeps are characterised by the migration of gas and oil from deep subsurface sources to the sediment-water interface^1, 2^. This seepage often contains gaseous short-chain alkanes, as well as heavier liquid alkanes and aromatic compounds^3^, that originate from deep thermogenic petroleum deposits. Migrated hydrocarbons can serve as an abundant source of carbon and energy for microorganisms in these ecosystems, either via their direct utilization or indirectly through metabolizing by-products of hydrocarbon biodegradation^4^. Multiple 16S rRNA gene surveys have revealed that cold seep sediments at or near the sediment-water interface host an extensive diversity of archaeal and bacterial lineages^5–9^. However, much less is known about metabolic versatility of this diverse microbiome, and how surface and subsurface populations are connected and differentiated in different redox zones within the sediment column^10, 11^. Most seep-associated microorganisms lack sequenced genomes, precluding meaningful predictions of relationships between microbial lineages and their biogeochemical functions^4, 8, 10–13^.

Geochemical studies have provided evidence that microorganisms in deep seafloor sediments, including cold seeps, mediate anaerobic hydrocarbon oxidation^2, 3^. A range of efforts have been undertaken to enrich and isolate anaerobic hydrocarbon-oxidizing microorganisms from cold seep sediments and other ecosystems rich in hydrocarbons (e.g. marine hydrothermal vents)^5–9, 14^. Numerous studies have focused on anaerobic oxidation of methane, since methane generally is the dominant hydrocarbon in cold seep fluids. This process is mediated by anaerobic methanotrophic archaea (ANME) through the reverse methanogenesis pathway, typically in syntrophy with bacteria that can reduce electron acceptors such as sulfate, nitrate, and metal oxides^8, 15, 16^. Investigations of enrichment cultures have also revealed anaerobic bacterial or archaeal oxidation of non-methane alkanes and aromatic hydrocarbons, including ethane (e.g. *Ca.* Argoarchaeum)^17^, *n*-butane (e.g. *Ca.* Syntrophoarchaeum and *Desulfobacteraceae* BuS5)^9, 18^, dodecane (e.g. *Desulfosarcina/Desulfococcus* clade)^6^, and naphthalene (e.g. deltaproteobacterial strain NaphS2)^19, 20^. In parallel, culture-independent approaches have provided a holistic *in situ* perspective on hydrocarbon biodegradation, with most studies focused on hydrothermally influenced sediments that are rich in hydrocarbons^21–24^. These studies have provided insights into the phylogenetic diversity and functional capabilities of potential hydrocarbon-degrading microorganisms, as well as their potential metabolic interactions with other community members. In addition to hydrocarbons, microbial life in deep-sea sediments is also supported by other various chemicals, including necromass-derived compounds and inorganic electron donors^4, 25^.

At present, in contrast to studies of deep-sea hydrothermal sediments^21–23^, there have been fewer reports on the metabolism of hydrocarbons and other compounds in cold seep sediments, especially in the deep sea^4^. Single-gene surveys, for example investigating the diversity of genes encoding enzymes for alkane or aromatic compound activation via addition to fumarate^3, 11, 26^, have increased our knowledge about the phylogenetic diversity of hydrocarbon degraders. However, more integrative approaches connecting geochemical processes to underlying microbial communities at cold seeps are lacking. In this study, we combine geochemical and metabolomic analyses with gene- and genome-centric metagenomics to understand the communities and processes responsible for anaerobic oxidation of different hydrocarbons, as well as other associated metabolisms, at a deep sea cold seep. The Scotian Basin is at the volcanic and non-volcanic transition continental margin, extending over an area of ∼260,000 km^2^ in the northwest Atlantic Ocean, offshore Nova Scotia in eastern Canada. Based on satellite and seismic reflection data, this area shows strong evidence for seepage of thermogenic hydrocarbons with occurrences of high pressure diapirs, polygonal faults, pockmarks, and gas chimneys^27^. Through this work, we provide strong evidence supporting that (i) hydrocarbons are thermogenic and experience biodegradation upon migration up into surface sediment layers; (ii) these processes are actively performed in the cold deep sea by bacteria and archaea through diverse biochemical mechanisms; and (iii) biodegradation is central to ecosystem productivity and supports various ecological interactions within the microbial communities.

## Results

### Thermogenic hydrocarbons migrate up the seep and are subject to biodegradation

At a water depth of 2306 m, a 3.44-meter-long piston core was retrieved from the Scotian slope, off the coast of Eastern Canada **(Supplementary Figure 1)**^27^. In the bottom of the core, at 332-344 cm below the seafloor (cmbsf), gas hydrates were observed in frozen crystalized form and numerous gas bubbles escaped during retrieval. A strong sulfide odor was also detected during core retrieval and processing. Molecular and isotopic composition of two headspace gas samples subsampled from the sediments adjacent to the gas hydrates (332-337 and 337-344 cmbsf) detected 7,446 and 4,029 ppm of total hydrocarbon gases (THG), respectively, made up of primarily methane (85% and 79%) as well as considerable proportions of C_2_-C_4_ gases (3.22% and 6.54%) **(Table 1)**. In order to assess the origin of the hydrocarbon gases, their molecular and isotopic compositions were compared to recently revised genetic diagrams^28^, focusing on δ^13^C-C_1_, C_1_/(C_2_+C_3_), δ^2^H-C_1_, δ^13^C-C_1_ and δ^13^C-CO_2_. All measured geochemical parameters are within the range defined for gases of thermogenic origin indicating that they migrated upward from a mature petroleum source rock **(Table 1)**. Ethane and propane were ^13^C-enriched compared with methane, likely reflecting the addition of some biogenic methane to the migrating thermogenic gas, as well as biodegradation of ethane and propane^28^. Ratios of *iso*-butane to *n*-butane were 1.6 to 1.8 and suggest preferential consumption of more labile *n-*alkanes^3^.

**Table 1.** Molecular and isotopic compositions of two gas samples sourced from sediments subsampled from the core regions nearest gas hydrates.

| Depth (cmbsf) |  | 332-337 | 337-344 |
| --- | --- | --- | --- |
| Features |  | Bubbling gas | Gas hydrate |
| Gas compositions<br>(% THG) | Methane | 85.10 | 79.00 |
|  | Ethane | 2.00 | 1.38 |
|  | Propane | 3.10 | 1.25 |
|  | *iso*-butane | 0.93 | 0.36 |
|  | *n*-butane | 0.51 | 0.23 |
| | $\Sigma C_2-C_4$ | 6.54 | 3.22 |
|  | THG (ppm) | 7446 | 4029 |
| | $C_1/(C_2+C_3)$ | 16.7 | 30.0 |
| Isotopic signatures<br>(‰) | $C_1 \delta^{13}C$ | -42.2 | -49.0 |
| | $C_1 \delta^2H$ | -169 | -177 |
| | $C_2 \delta^{13}C$ | -24 | -26 |
| | $C_3 \delta^{13}C$ | -21.5 | -22.3 |
| | *i*-$C_4 \delta^{13}C$ | -22.6 | ND |
| | *n*-$C_4 \delta^{13}C$ | -21.6 | ND |
| | $CO_2 \delta^{13}C$ | -6.3 | -7.7 |

Abbreviations: $C_1$ , methane; $C_2$ , ethane; $C_3$ , propane; $C_4$ , butane. THG, total hydrocarbon gases. $C_1/(C_2+C_3)$ , molecular ratios of methane to ethane and propane. $\delta^{13}C$ and $\delta^2H$ , stable carbon and hydrogen isotope composition. ND, not determined.

Sediments from four additional depths were analyzed for extractable organic matter (EOM, i.e., C_12+_ hydrocarbons), showing high yields (104-361 mg/kg rock) comprising saturated hydrocarbons (25-52%), aromatic hydrocarbons (10-14%), and other components **(Supplementary Table 1)**. Further gas chromatographic analysis of these oil-laden sediments revealed large unresolved complex mixture (UCM) humps in the C_13_-C_20_ *n-*alkane elution range **(Supplementary Figure 2)**, indicative of oil biodegradation^3^. Pristane and phytane, widely used internal conserved markers for oil biodegradation^29^, were more abundant than C_17_ and C_18_ n-alkanes **(Supplementary Figure 2)**, suggesting preferential biodegradation of *n-*alkanes. Consistent with this, carbon dioxide was isotopically heavy in these sediments **(Table 1)** and clearly associated with secondary microbial degradation^28^.

### Microbial communities are phylogenetically diverse and strongly stratified across biogeochemical zones

Deep shotgun metagenome sequencing was performed for sediments corresponding to four biogeochemical redox zones based on pore water sulfate concentrations and other geochemical data **(Table 2, Supplementary Figure 3 and Supplementary Note 1)**. Alpha diversity **(Table 2)** was calculated using single-copy marker genes from the metagenomic datasets^30^, and cell density was estimated via quantitative PCR of bacterial and archaeal 16S rRNA genes **(Table 2)**. Differences in diversity and cell density were related to the availability of migrated thermogenic hydrocarbons and sulfate **(Table 1, Supplementary Table 1 and Supplementary Figure 3)**. Overall, the surface sediment (mixing zone) and underlying sediment (sulfate reduction zone) harboured the most diverse communities (Shannon index = 6.70 and 5.58) and highest bacterial cell density (2.64 and 2.22 × 10^9^ 16S rRNA genes g^−1^). In contrast, the sediment at 60 cmbsf (sulfate methane transition zone) harboured the most distinct communities **(Figure 1a)**, the lowest microbial diversity (3.60), and the highest archaeal cell density (1.23 × 10^9^ 16S rRNA genes g^−1^). Deeper sediments (100 to 250 cmbsf; methanogenic zone) were more compositionally similar and harboured moderately to highly diverse and abundant communities **(Table 2 and Figure 1a**).

**Figure 1.**
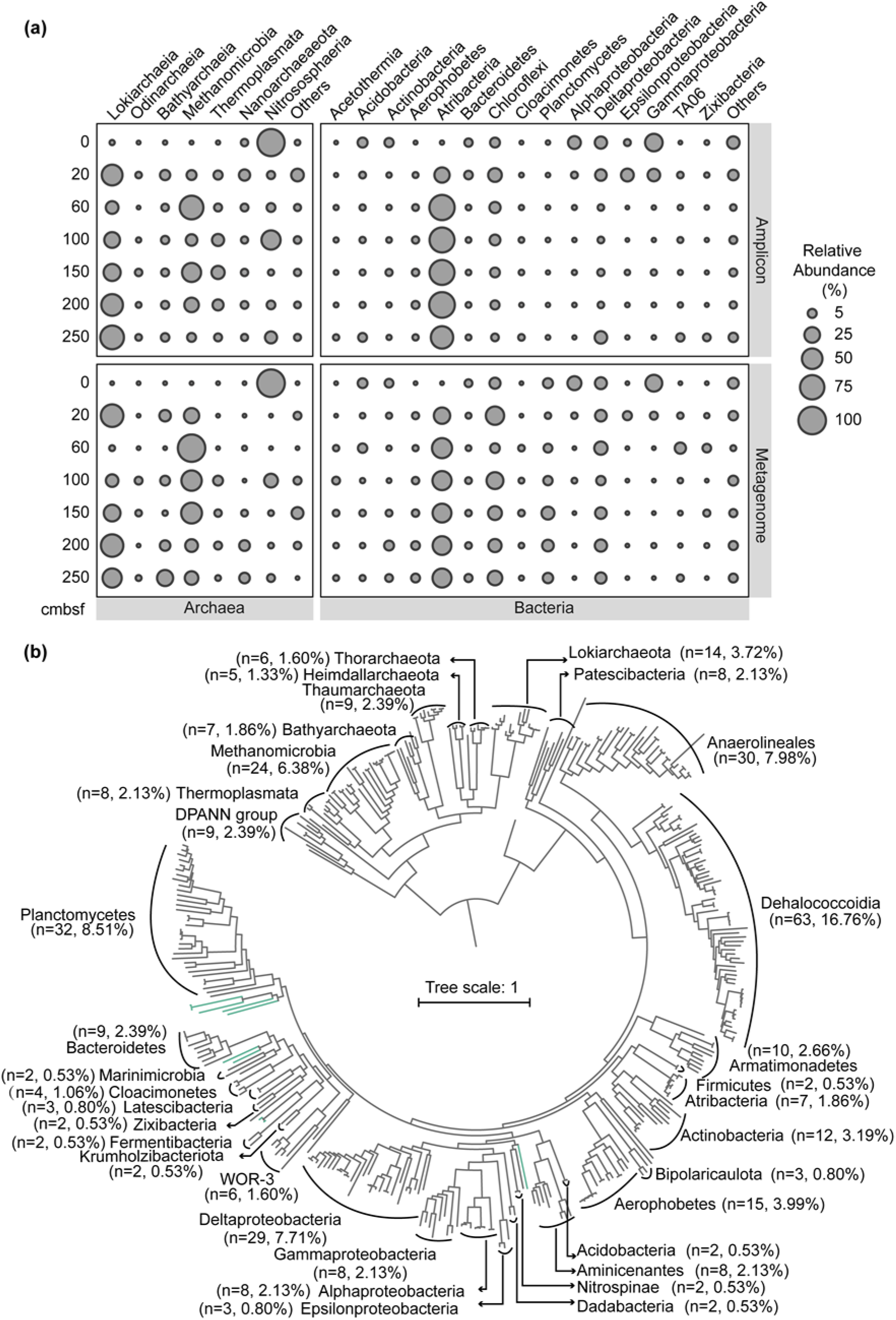
Gene- and genome-resolved view of the microbial communities. (a) Distribution of bacterial and archaeal taxa at different sediment depths. Top panels: relative abundance based on 16S rRNA gene amplicon analysis; bottom panels: reconstruction of full-length 16S rRNA genes from the metagenomes. Archaeal groups are shown on the left and bacterial groups are shown on the right. (b) Phylogenetic placement of 376 reconstructed metagenome-assembled genomes. A maximum-likelihood phylogenomic tree was built based on concatenated amino acid sequences of 43 conserved single-copy genes using RAxML with the PROTGAMMALG model. The scale bar represents 1 amino acid substitution per sequence position. Numbers in parentheses show total MAGs recovered. Phyla with only one detected MAG are not labelled. Ten MAGs corresponded to unclassified phylum-level lineages (green branches; unlabelled). The full tree file with references is available in **Supplementary Data 1**.

**Table 2.**
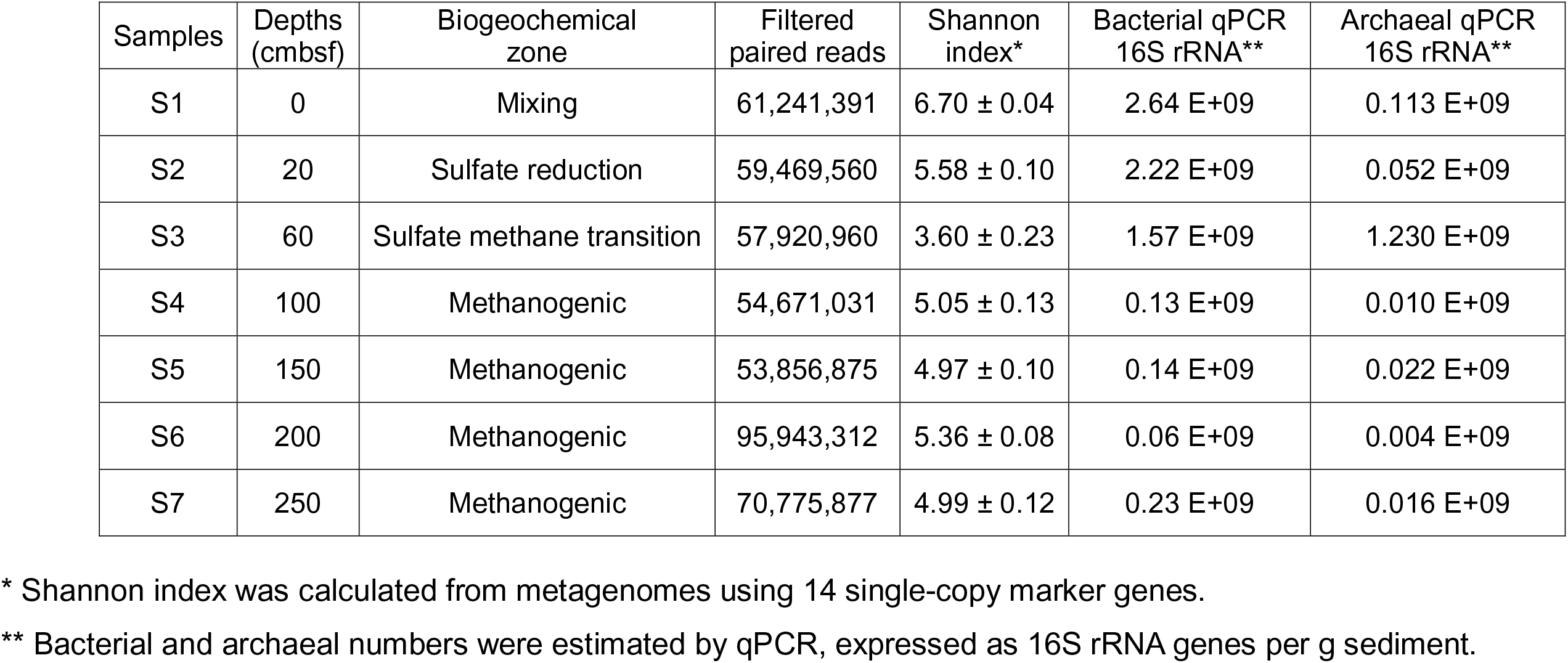
Key features of sediment samples used for microbiological analyses.

For taxonomic profiling, the phyloFlash pipeline^31^ was applied to reconstruct 16S rRNA gene sequences from metagenome raw reads for each sediment depth **(Figure 1a**). Dominant bacterial lineages in surface sediments were *Gammaproteobacteria* (30%) and *Alphaproteobacteria* (19%), whereas Atribacteria (28-47%), Chloroflexi (6-37%), *Deltaproteobacteria* (6-16%), and Planctomycetes (1-15%) predominated at and below 20 cmbsf **(Figure 1a**). As globally abundant groups in subseafloor sediments at ocean margins^32, 33^, Atribacteria and Chloroflexi accounted for most of the bacterial communities sampled. Deeper layers also contained candidate phyla largely absent from shallower layers, e.g. TA06, Acetothermia, Zixibacteria, and Aerophobetes. Thaumarchaeota (mainly class *Nitrososphaeria*), the only dominant archaea found in surface sediment, were in low abundance in subsurface sediments **(Figure 1a**). *Methanomicrobia* (Euryarchaeota) and Lokiarchaeota (Asgard group) predominated at and below 20 cmbsf, with the former comprising 99% of archaea at 60 cmbsf, consistent with this being the sulfate methane transition zone **(Supplementary Figure 3)**. *Ca.* Bathyarchaeota, *Ca.* Nanoarchaeota, *Thermoplasmata*, and *Ca.* Odinarchaeota were also abundant in certain layers. Taxonomic profiles produced by 16S rRNA gene amplicon sequencing were broadly similar to metagenomic profiling, but with differences in the relative abundances of specific groups (e.g. Chloroflexi and *Nitrososphaeria*) **(Figure 1a)**. Together, both methods show that microbial communities throughout the sediment column are diverse and consist of mostly uncultured taxonomic groups.

Metagenomes were assembled for sediments from individual depths, and a co-assembly was performed by combining metagenomes from all depths^34^ **(Supplementary Tables 2 and 3)**. Binning of derived assemblies was based on tetranucleotide frequencies and coverage profiles using several algorithms^35^. This analysis yielded 376 unique metagenome-assembled genomes (MAGs; 293 bacterial and 83 archaeal) with >50% completeness and <10% contamination based on CheckM analysis^36^. Recovered MAGs spanned 43 different phyla, most of which are poorly characterized without cultured representatives **(Figure 1b, Supplementary Table 2 and Supplementary Data 1)**. Ten of these genomes could not be classified due to lack of reference genomes and, based on their phylogeny, may belong to five new candidate phyla **(Figure 1b and Supplementary Data 1)**. Overall the 376 MAGs captured the prevalent bacterial and archaeal lineages revealed by 16S rRNA gene analysis **(Figure 1a)**. The MAGs with the highest relative abundance belong to the ANME-1 lineage, and together made up >40% of the microbial community based on read coverage in the sulfate methane transition zone at 60 cmbsf **(Supplementary Table 2)**. Among ANME-1 genomes at this depth, the *in situ* replication rate inferred from iRep was relatively low (e.g. iRep score of 1.38 for S3_bin12)^37^, suggesting slow replication despite high relative abundance (e.g. 4.69% for S3_bin12) **(Supplementary Table 2)**. MAGs corresponding to Atribacteria, Lokiarchaeota, and Chloroflexi were also differentially enriched with depth, consistent with their dominance in these sediments and their occurrence in the deep marine biosphere more generally^32, 33^. Among these groups, 22 Chloroflexi MAGs were determined to have high genome replication rates, with a maximum iRep value of 4.0 for *Dehalococcoidia* S5_bin22 in the methanogenic zone. This suggests several replication forks at the time of sampling, providing strong evidence for activity by *Dehalococcoidia*. Most other MAGs were much less abundant (e.g. <0.1%). Some nevertheless had large iRep values, for example *Ca.* Thorarchaeota S7_bin1 from the Asgard group (iRep = 2.1) and *Desulfobacterales* S3_bin8 (iRep = 6.1), with relative abundances of 0.1% at 200 cmbsf and 0.28% at 150 cmbsf, respectively **(Supplementary Table 2)**.

### Diverse Euryarchaeota potentially mediate anaerobic oxidation of methane and other short chain alkanes

Archaea activate short-chain alkanes (methane, ethane, propane, and butane) for anaerobic degradation using methyl/alkyl-coenzyme M reductases^17^. Sequences encoding the catalytic subunit of this enzyme (*mcrA*) were detected in metagenomes at all sediment depths except 0 cmbsf, with the highest abundance found at the sulfate methane transition zone **(Figure 2a)**. A total of 20 MAGs within Euryarchaeota harbored *mcrA* sequences (**Figure 2b and Supplementary Table 4)**. Genome trees showed these microorganisms belonged to representatives of three families of hydrogenotrophic methanogens (*Methanomicrobiaceae*, *Methanosarcinaceae* and *Methanosaetaceae*)^38^, two clusters of anaerobic methanotrophs (ANME-1 and ANME-2)^39^, and two lineages that have been shown to catalyse non-methane alkane oxidation (the GOM-Arc1 lineage, and a novel sister lineage to *Ca.* Syntrophoarchaeum)^17, 22^ **(Figure 3a)**. In agreement with this, phylogenetic analysis of *mcrA* sequences from these MAGs resolved three major groups **(Figure 3b)**: the canonical group clustering with methanogens and methanotrophs (ANME-1 and ANME-2) and two divergent groups clustering with *Ca.* Syntrophoarchaeum^18^ and GOM-Arc1 (e.g. *Ca.* Argoarchaeum)^17^. These phylogenetic relationships, together with presence of the mixture of alkane gases and their carbon isotope ratios **(Table 1)** suggest that anaerobic methane- and multi-carbon alkane-metabolizing archaea coexist in these cold seep sediments. These findings mirror those recently reported in the hot Guaymas Basin hydrothermal sediments^40^ and thus reveal that recently discovered C_2+_ short chain alkane metabolisms are not peculiar to hot hydrothermal settings and are likely much more widespread throughout the deep seabed at cold seeps. A notable difference from other cold seep sediments is that this Scotian Basin site is dominated by ANME-1 rather than ANME-2 lineages^10, 11^, with three distinct ANME-1 groups (S3_bin4, Co_bin174 and S3_bin12) being particularly abundant (27.47%, 7.33% and 4.69%, respectively) at the sulfate methane transition zone **(Supplementary Table 2)**, an observation supported by 16S rRNA gene amplicon sequencing **(Supplementary Table 5)**.

**Figure 2.**
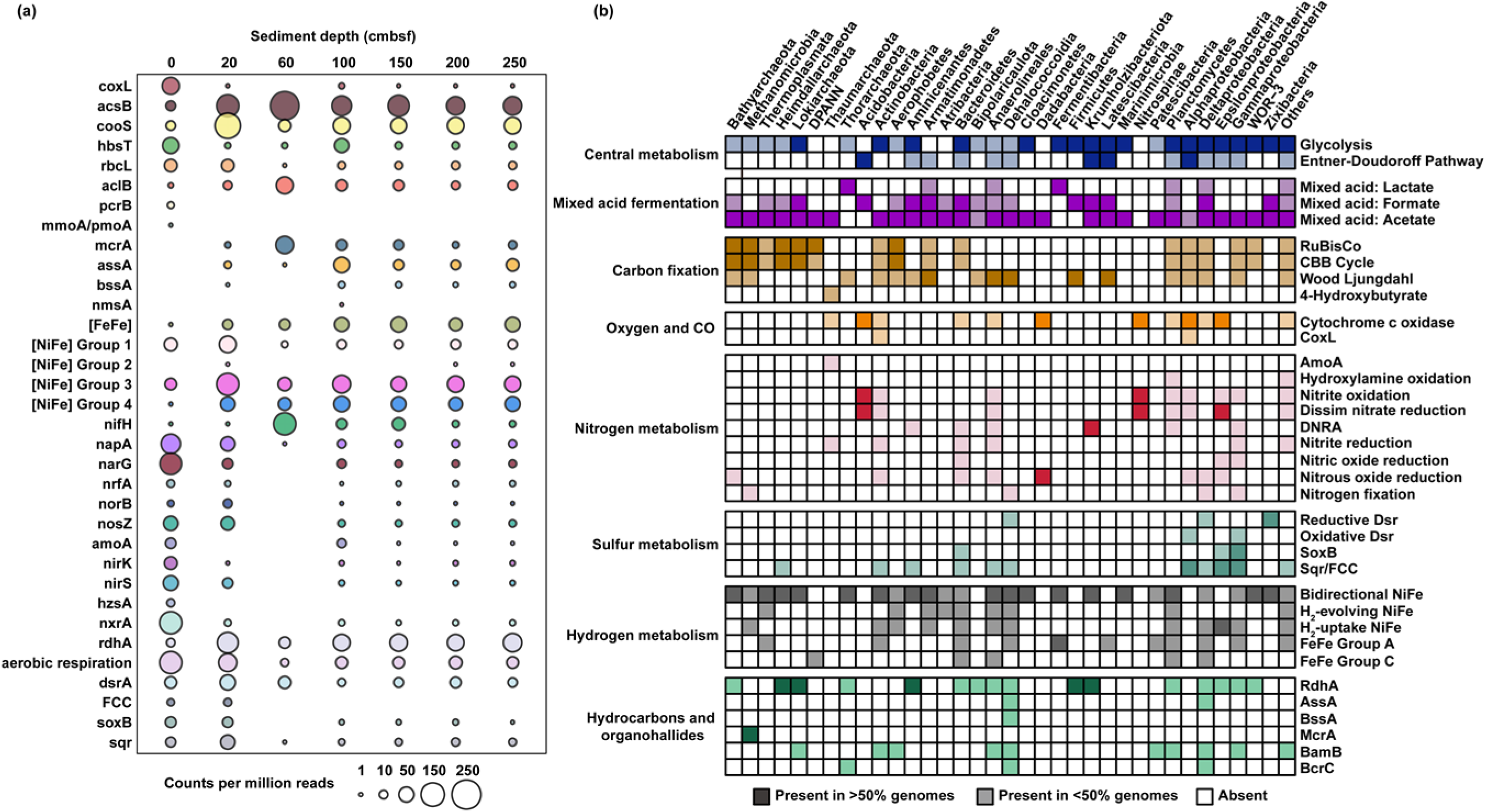
Occurrence of core metabolic genes or functions in unassembled quality-filtered reads and phylogenetic clusters of MAGs. (a) Depth-differentiated metabolic capacity based on normalized short-read counts of genes encoding the catalytic subunits of key metabolic enzymes in unassembled quality-filtered reads. (b) Core metabolic genes or pathways and their occurrence (in %) in MAGs grouped into 35 different phylogenetic clusters. Metabolic categories (from top to bottom): central metabolism, mixed acid fermentation, carbon fixation, oxygen and carbon monoxide metabolism, nitrogen metabolism, sulfur metabolism, hydrogen metabolism, reductive dehalogenation, and hydrocarbon degradation. Phyla with only one representative MAG in the dataset (n=10) and MAGs that could not be classified (n=10) are grouped together (Others; far right). Shaded colors: gene present in 1-50% of genomes/phylogenetic cluster. Solid colors: gene present in 50-100% of genomes/cluster. Complete lists of metabolic genes or pathways can be found in **Supplementary Tables 4 and 6-12**.

**Figure 3.**
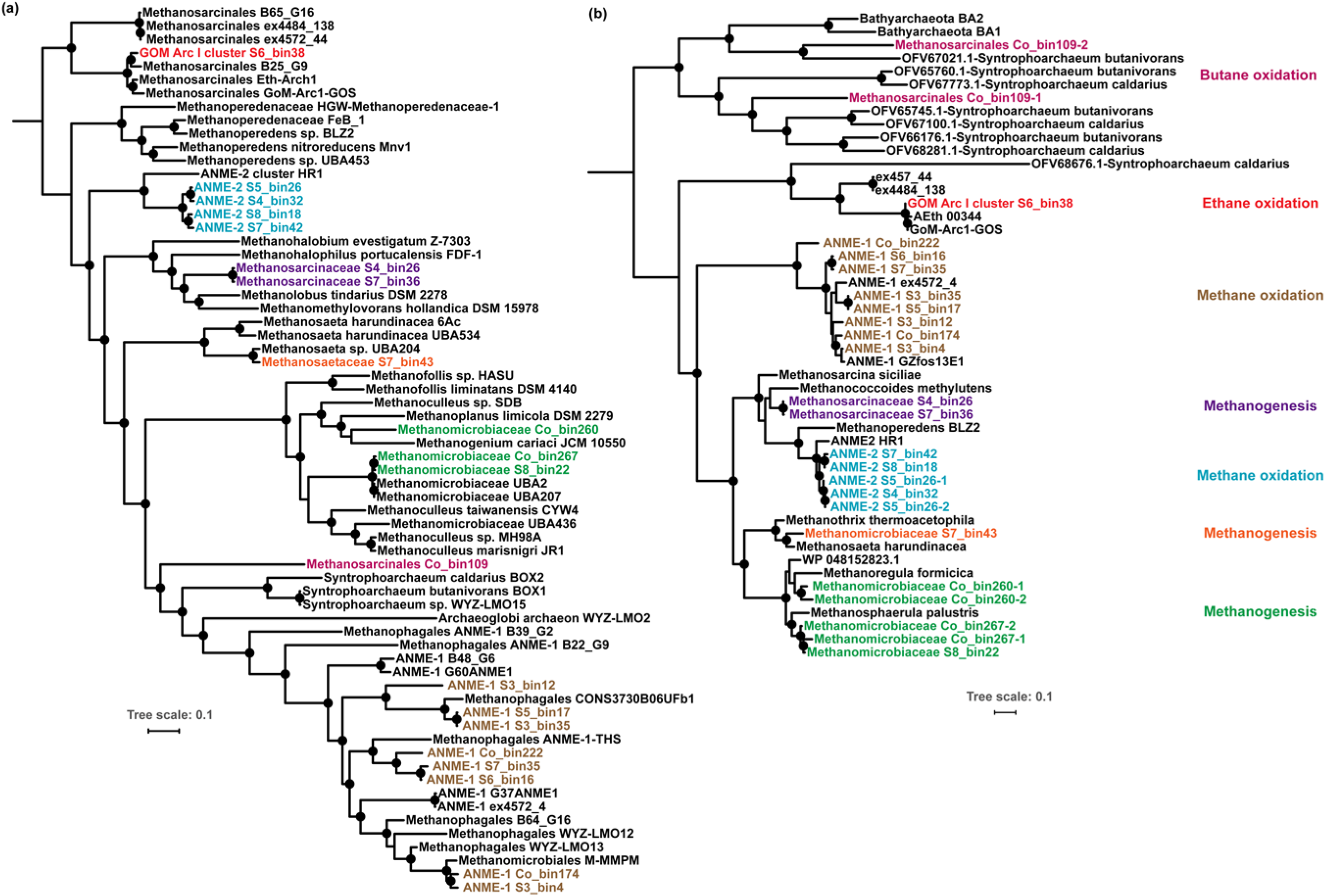
Maximum likelihood phylogenetic trees of MAGs and the detected methyl-Coenzyme M reductase (McrA) sequences. (a) Phylogenomic tree constructed based on alignments of 43 conserved protein sequences from *Methanomicrobia* MAGs; (b) phylogenetic tree constructed based on alignments of amino acid sequences of *mcrA* genes. Bootstrap values of >70% are indicated with black circles. MAGs and McrA sequences corresponding to the same alkane metabolisms are highlighted in the same color in the two trees. Scale bars correspond to per cent average amino acid substitution over the alignment, for both trees. Sequences for amino acids used to construct McrA trees can be found in **Supplementary Data 2**.

To gain more detailed insights into anaerobic short chain alkane degradation in this non-hydrothermal setting, metabolic pathways involved in the oxidation of methane and non-methane gaseous alkanes were reconstructed^17^ **(Figure 4)**. Twelve MAGs harbour canonical *mcrA* genes that cluster with ANME-1 and ANME-2 methanotrophs, along with with *fwd*, *ftr*, *mch*, *mtd*, *mer/metF/fae-hps* and *mtr* that mediate subsequent steps in the tetrahydromethanopterin-dependent ‘reverse methanogenesis’ pathway^10, 41^ for oxidation of methyl-CoM to CO_2_ **(Figure 4a and Supplementary Table 6)**. Some MAGs lack specific genes in this pathway, likely reflecting variability in genome completeness (61% to 96%). Based on genome and gene trees, S6_bin38 is closely related to putative ethane oxidizers within the GoM-Arc1 group^21, 22, 38^, including the verified anaerobic ethane oxidizer *Ca.* Argoarchaeum ethanivorans Eth-Arch1^17^ **(Figure 3)**. Like other GoM-Arc1 genomes, S6_bin38 encodes methyltransferases that potentially transfer the thioether (ethyl-CoM) derived from ethane activation to a thioester (acetyl-CoA), as well as enzymes to mediate acetyl-CoA cleavage (acetyl-CoA decarbonylase/synthase, ACDS) and stepwise dehydrogenation of the derived C_1_ units (oxidative Wood-Ljungdahl pathway) **(Figure 4b and Supplementary Table 6)**. S6_bin38, like other members of the GoM-Arc1 group, lacks the β-oxidation pathway which is unnecessary for anaerobic ethane oxidation^17^. Taken together, these reconstructions suggest that S6_bin38 represents an organism capable of oxidizing ethane in cold deep sea sediment.

**Figure 4.**
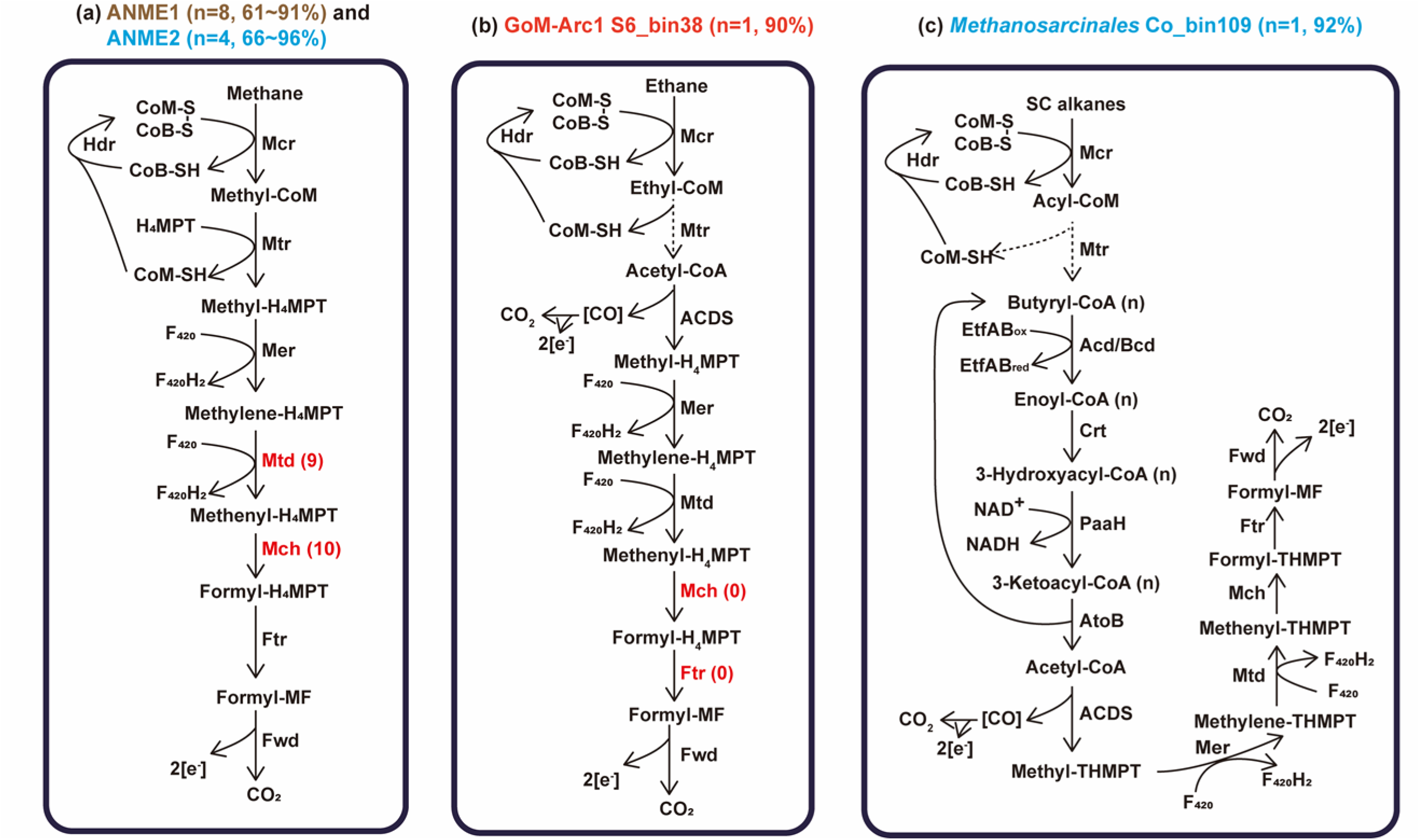
Predicted metabolic models for anaerobic oxidation of gaseous alkanes based on archaeal MAGs harboring *mcrA* genes. (a) Archaeal oxidation of methane; (b) archaeal oxidation of ethane; and (c) archaeal oxidation of butane. Red font indicates that not all MAGs retrieved encode the enzyme (numbers of MAGs encoding the enzyme are indicated in parentheses). Dashed lines indicate steps catalyzed by unconfirmed enzymes. SC alkanes, short chain alkanes. Percentages above each panel indicate the completeness of the corresponding MAGs estimated by CheckM. Details on the annotation of the enzymes are presented in **Supplementary Table 6**.

*Methanosarcinales* Co_bin109 likely has potential to oxidize butane and fatty acids that were present in the sediments (**Table 1 and Figure 5)**. This genome contains two *mcrA* genes that cluster with high bootstrap support to the divergent alkyl-CoM reductases from *Ca.* Syntrophoarchaeum **(Figure 3b)** that is capable of anaerobic degradation of butane and possibly propane^18^. Also encoded in the genome are heterodisulfide reductase subunits (*hdrABC*) to reoxidize cofactors^23^, methyltransferases that potentially convert butyl-thioether to the butyryl-thioester, and the β-oxidation pathway to enable complete oxidation of the butyryl-thioester **(Figure 4c and Supplementary Table 6)**. The presence of short chain acyl-CoA dehydrogenase (*acd*), butyryl-CoA dehydrogenase (*bcd*), and long-chain acyl-CoA synthetase (*fadD*) may allow *Methanosarcinales* Co_bin109 to oxidize long chain fatty acids; this is similar to the basal *Archaeoglobi* lineage *Ca.* Polytropus marinifundus, which encodes two divergent McrA related to those found in *Ca.* Bathyarchaeota and *Ca.* Syntrophoarchaeum^42^. Consistent with this, Co_bin109 encodes a Wood-Ljungdahl pathway (including carbon monoxide dehydrogenase / acetyl-CoA synthase complex) for complete fatty acid oxidation **(Supplementary Table 6)**.

**Figure 5.**
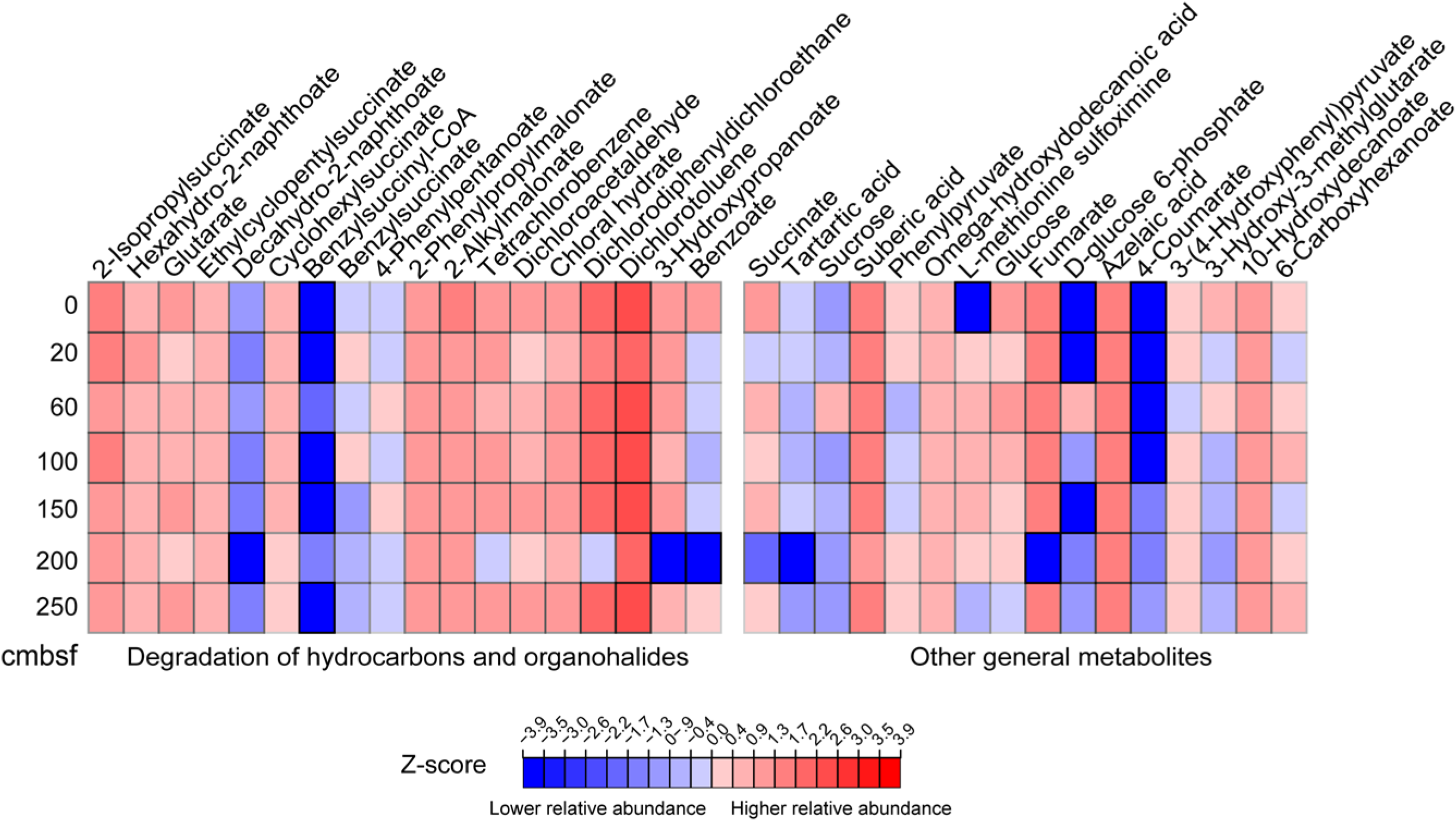
Heatmap of metabolites identified in sediment pore water from down core sediment subsamples. Metabolite levels were measured using HPLC Orbitrap mass spectrometry and are expressed as cumulative sum logarithmically normalized peak areas of sample replicates (n = 3). Compound names are listed above the heatmap.

### Members of Chloroflexi and *Deltaproteobacteria* potentially degrade liquid alkanes and aromatic hydrocarbons

Other community members are predicted to degrade aromatic hydrocarbons and *n-* alkanes detected in the sediments **(Table 1 and Supplementary Table 1)** via hydrocarbon addition to fumarate^43^. Nine MAGs assigned to *Dehalococcoidia*, *Desulfobacterales* and *Syntrophobacterales* encode alkylsuccinate synthase (*assA*) known to mediate *n-*alkane activation **(Figure 6a; Supplementary Tables 2 and 7)**. Phylogenetic analysis confirms that these *assA* genes cluster closely with those of experimentally validated alkane oxidizers *Desulfatibacillum aliphaticivorans* DSM 15576 and *Desulfosarcina* sp. BuS5 within Deltaproteobacteria^8, 44^ **(Supplementary Figure 4)**. In most of these MAGs, more than one *assA* sequence variant was identified, suggesting some bacteria may activate multiple substrates by this mechanism^45, 46^. In addition to *assA* sequences, *Dehalococcoidia* Co_bin57 and Co_bin289 also encode related glycyl-radical enzymes clustering with benzylsuccinate synthases (*bssA*) and naphthylmethylsuccinate synthases (*nmsA*) **(Supplementary Figure 4 and Supplementary Table 7)**, known to mediate the anaerobic degradation of toluene or similar aromatic compounds. Metabolomic analysis detected four succinic acid conjugates involved in hydrocarbon activation, including conjugates of both toluene and propane **(Figures 5 and 6)**. Most of the MAGs encoding *assA* or *bssA* genes (except *Dehalococcoidia* Co_bin57) also encode the required genes to further process the alkyl-/arylalkylsuccinate compounds, convert them to acetyl-CoA through the β-oxidation pathway, and regenerate fumarate through the methylmalonyl-CoA pathway **(Figure 6a and 6c; Supplementary Table 7)**. Accordingly, succinate and fumarate were also detected in the sediments **(Figure 5)**.

**Figure 6.**
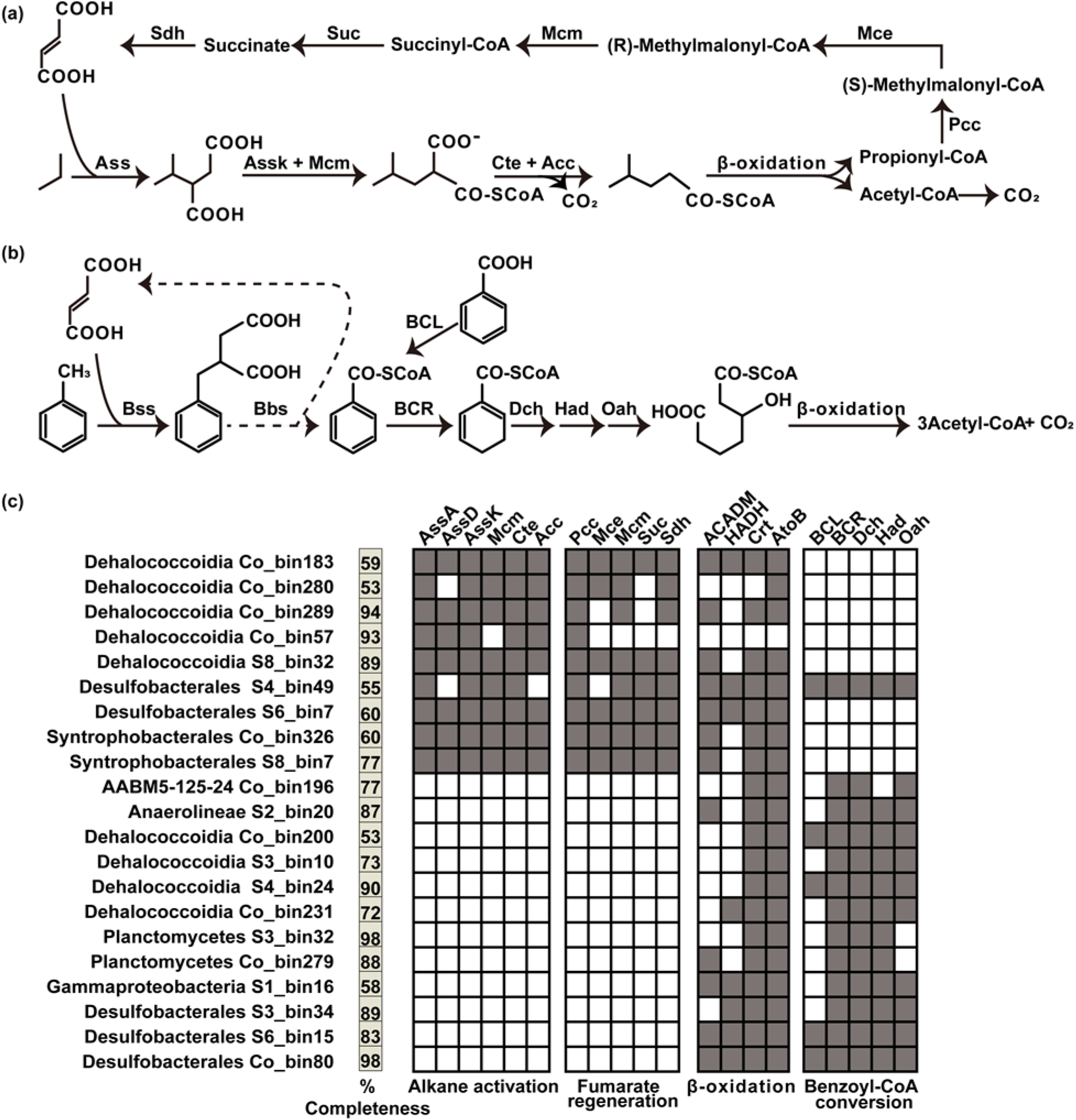
Predicted metabolic models for anaerobic degradation of alkanes and benzoate based on bacterial MAGs harboring *assA* and *bcr* genes. Proposed metabolic pathways for anaerobic degradation of (a) propane and (b) toluene via addition to fumarate. Propane and toluene here serve as examples for illustrating anaerobic degradation of alkane and aromatic hydrocarbons more generally. (c) Occurrence of protein encoding genes involved in these pathways in bacterial MAGs, as a presence/absence (filled/white squares) matrix. Genome completeness (%) for each MAG as estimated by CheckM is indicated. Details on the annotation of the enzymes are presented in **Supplementary Tables 7 and 8**.

Anaerobic hydrocarbon degradation depends on subsequent oxidation of acetyl-CoA. This can be achieved through a single-cell process (e.g. coupled to sulfate respiration) or via syntrophic interaction with another cell (e.g. with methanogens)^8, 44^. Like the related deltaproteobacterial isolate NaphS2, *Desulfobacterales* MAGs S4_bin49 and S6_bin7 contain genes for both the Wood-Ljungdahl pathway and dissimilatory sulfate reduction, suggesting the same organism can couple alkane mineralisation directly to sulfate reduction **(Supplementary Table 7)**. By contrast, other MAGs containing genes for hydrocarbon addition to fumarate, including *Dehalococcoidia* and *Syntrophobacterales*, apparently lack terminal reductases or complete tricarboxylic acid cycles **(Supplementary Table 7)**. These organisms may be obligate fermenters dependent on syntrophy with respect to the oxidation of *n*-alkanes, as further evidenced by identification of genes for mixed-acid fermentation and hydrogen production in these genomes **(Supplementary Tables 4 and 7)** and as seen in closely related organisms from both lineages^45^. Several MAGs found at the same depths were identified as methanogens **(Figure 3b and Supplementary Table 4)**, thus syntrophic alkane mineralization via hydrogenotrophic or acetoclastic methanogenesis is another route for hydrocarbon biodegradation in this environment. Accordingly, potential alkane degraders like *Syntrophobacterales* S8_bin7 were mainly found in the deeper methanogenic zone, co-occurring with methanogens such as *Methanomicrobiaceae* S8_bin22 **(Figure 3b and Supplementary Table 2)**.

Detection of *bssA* and *nmsA* suggest these cold seep microbial communities are also capable of utilizing aromatic hydrocarbons that were detected in the GC-MS analysis **(Figure 2a; Supplementary Table 1 and Supplementary Figure 4).** Metabolites produced after initial activation normally channel into the central benzoyl-CoA degradation pathway **(Figure 6b)**. Benzoyl-CoA, as a universal biomarker for anaerobic degradation of aromatic compounds^47^, is reduced by benzoyl-CoA reductases of the ATP-dependent class I pathway (*bcr* genes; e.g., in *Thauera aromatica*) or ATP-independent class II pathway (*bam* genes; e.g., in sulfate reducers). These genes were detected in 13 bacterial MAGs, including *Dehalococcoidia*, *Anaerolineae*, *Deltaproteobacteria*, Planctomycetes, *Gammaproteobacteria*, and candidate phylum AABM5-125-24. Genes for further processing these compounds to 3-hydroxypimelyl-CoA (i.e. *oah*, *dch* and *had*) and acetyl-CoA (β-oxidation pathway) were also detected **(Figure 6c and Supplementary Table 8)**. Various compounds related to benzoyl-CoA degradation pathway were also detected, including benzoate and glutarate (**Figure 5**).

### Metabolomic profiling suggests the community is supported by diverse energy conservation and carbon acquisition strategies

To understand the wider metabolic capabilities of these cold seep microbial communities, metabolomic analysis **(Figure 5)** was combined with gene-centric analysis of quality-filtered reads **(Figure 2a)** and unbinned metagenome assemblies **(Supplementary Note 2 and Supplementary Figure 5)**, as well as genome-centric analysis of the 376 MAGs **(Figure 2b).** With respect to organotrophy, genes for the degradation of complex carbohydrates and peptides were prevalent. In total, ∼7454 potential carbohydrate-active enzymes (CAZymes) were detected, most notably glycoside hydrolases, carbohydrate esterases, and polysaccharide lyases^48^; CAZymes were numerous in bacterial lineages such as Planctomycetes, Bacteroidetes, and Armatimonadetes **(Supplementary Table 9)**. Annotated peptidases were more evenly distributed across bacteria and archaea, with the highest proportions found in genomes assigned to Patescibacteria, *Dehalococcoidia*, Aerophobetes, and Atribacteria **(Figure 2b and Supplementary Table 10)**. Consistent with these genomic observations, metabolites associated with carbohydrate and peptide hydrolysis were detected (e.g. glucose and sucrose) **(Figure 5)**. Many community members also possess gene sets to convert hydrolysis products to pyruvate through the Embden-Meyerhof-Parnas (185 MAGs) and Entner-Doudoroff (51 MAGs) pathways **(Figure 2b and Supplementary Table 4)**. Accordingly, both glucose 6-phosphate and pyruvate were detected in the sediment column metabolomes **(Figure 5)**. Most genomes also encoded the potential to ferment pyruvate to formate (119 MAGs), acetate (270 MAGs), lactate (14 MAGs), and ethanol (94 MAGs), as well as the potential for hydrogen production via abundant [FeFe]-hydrogenases associated with fermentation or group 3 [NiFe]-hydrogenases^49^ **(Figure 2a and 2b; Supplementary Figure 6 and Supplementary Table 4)**. The ability to degrade fatty acids and other organic acids via beta-oxidation was common to Lokiarchaeota, Chloroflexi, and Proteobacteria **(Supplementary Table 11)**, with associated metabolites also detected (e.g. 6-carboxyhexanoate, 10-hydroxydecanoate, and other carboxylic acids) **(Figure 5)**.

The most prevalent carbon fixation pathways in these sediments are the Calvin-Benson cycle (68 MAGs), Wood-Ljungdahl pathway (114 MAGs), and 4-hydroxybutyrate cycle (4 MAGs) **(Figure 2b and Supplementary Table 4)**, consistent with the abundance of corresponding key metabolic genes **(Figure 2a)**. The sediment microbial communities include various putative sulfide, thiosulfate, ammonia, nitrite, carbon monoxide, and hydrogen oxidizers **(Figure 2b and Supplementary Table 4)**, with provision of these compounds potentially via biological or geological processes. Canonical genes for sulfide oxidation (sulfide:quinone oxidoreductase and flavocytochrome *c* sulfide dehydrogenase) were found in 38 MAGs from diverse lineages (e.g. Proteobacteria, *Anaerolineales*, and Heimdallarchaeota) **(Supplementary Figures 7-8)**. The oxidative bacterial-type DsrAB-type dissimilatory sulfite reductases^50^ were detected in four genomes from *Alpha-* and *Gammaproteobacteria* potentially responsible for sulfide oxidation **(Supplementary Figure 9)**. Five MAGs from *Epsilon*- and *Gammaproteobacteria* encoded genes for the SoxB component of the periplasmic thiosulfate-oxidizing Sox enzyme complex **(Supplementary Figure 10).** Genes for ammonia oxidation to hydroxylamine (ammonia monooxygenase) were detected in four Thaumarchaeota genomes **(Supplementary Figure 11)**, while those associated with further oxidation of nitrite (nitrite oxidoreductase) were tentatively found in five MAGs affiliated to *Anaerolineales*, Acidobacteria, Actinobacteria, and Planctomycetes **(Supplementary Figure 12)**. Gene abundance profiles suggest that these nitrogen metabolisms mainly occur in 0-20 cmbsf surface sediments **(Figure 2a)**. Three MAGs belonging to Actinobacteria, Actinobacteria, and *Alphaproteobacteria* encoded carbon monoxide dehydrogenase large subunit (CoxL), a marker for carbon monoxide oxidation **(Supplementary Figure 13)**. Ten phyla (58 MAGs) encoded group 1 and group 2 [NiFe]-hydrogenases associated with hydrogenotrophic respiration, with certain lineages, including *Deltaproteobacteria*, Planctomycetes, *Dehalococcoidia*, Actinobacteria, and Aerophobetes, encoding subgroups (1a, 1b) that support hydrogenotrophic sulfate reduction and halorespiration^51, 52^ **(Figure 2b and Supplementary Figure 14; Supplementary Tables 4 and 12)**. Additionally, 197 MAGs from 35 phyla encode cofactor-coupled group 3 [NiFe]-hydrogenases (**Supplementary Figure 6)** and energy-converting group 4 [NiFe]-hydrogenases (**Supplementary Figure 15)**, both are which are known to be physiologically reversible^49^.

Metabolomics revealed organohalides as potential respiratory electron acceptors in the sediments, e.g. dichlorotoluene and dichlorodiphenyldichloroethane **(Figure 5)**. Accordingly, reductive dehalogenases were encoded in 54 MAGs from 13 phyla, including lineages such as Chloroflexi, Lokiarchaeota, Bathyarchaeota, and Proteobacteria **(Figure 2, Supplementary Figure 16 and Supplementary Table 12)**. Organisms encoding these enzymes may conserve energy by coupling oxidation of H_2_ and other electron donors to the reduction of organochloride compounds like the ones detected in the sediments **(Figure 5)**. Full gene sets for dissimilatory reduction of sulfate to sulfide (*sat*, *aprAB* and *dsrAB*) were found in 12 genomes from *Deltaproteobacteria* and Zixibacteria **(Figure 2b, Supplementary Figure 9 and Supplementary Table 4)**. Thirty bacterial members of these sediment communities are predicted to use nitrate or nitrite as electron acceptors, though none encoded all of the genes necessary for complete denitrification or dissimilatory nitrate reduction to ammonia (**Supplementary Table 4)**. Terminal oxidases were also detected, but in line with oxygen only permeating the upper centimeters of marine sediments, these genes were mainly found in lineages such as Proteobacteria and Thaumarchaeota that dominated the upper sediment layers **(Figure 2a and 2b; Supplementary Table 4)**.

## Discussion

This work provides evidence that thermogenic short-chain alkane gases, larger alkanes, and aromatic compounds are biodegraded in Scotian Basin cold seep sediments in the NW Atlantic deep sea. Sequencing DNA extracted from different sediment depths obtained via piston coring in >2 km deep water enabled associating petroleum geochemistry with microbial populations and metabolites. We also demonstrate that archaea capable of oxidizing methane and short-chain hydrocarbons co-exist with bacteria capable of degrading larger aliphatic and aromatic compounds. These findings reveal that anaerobic hydrocarbon-degrading microorganisms are more phylogenetically and metabolically diverse, and geographically widespread, than previously known^5, 8, 17, 53^.

The capacity for anaerobic degradation of hydrocarbons was detected in 44 out of 367 MAGs, including members encoding genes for C_2+_ hydrocarbon oxidation that belong to the phyla Euryarchaeota and Chloroflexi, in addition to well-known deltaproteobacterial lineages^21, 22^. Distributions of these organisms span from the near-surface sediment horizon to the deeper zones where sulfate is depleted, with overall relative abundance ranging from 1% to 42% at different depths. Identification of succinate derivatives^54^ (e.g. 2-isopropylsuccinate and benzylsuccinate) in pore water and determination of carbon stable isotopic signatures for hydrocarbon gases (e.g. ethane and propane) and carbon dioxide^28^ provide direct evidence of biodegradation actively occurring in these deep sea sediments. Phylogenetic analysis and pathway reconstruction shed new light on the different enzymes responsible for methane, ethane, and butane oxidation, relative to previous studies of anaerobic hydrocarbon degradation in cold seep sediments^5, 8, 17, 53^. Notably, the recently described process of multi-carbon alkane oxidation by archaea co-occurs with anaerobic methane oxidation by ANME archaea. Previous observations of these phenomena have mainly featured hydrothermal vent environments^15, 22, 38^ whereas this deep sea setting experiences permanently low temperatures. Methane oxidizers are most abundant, in agreement with the high concentration of methane; however, by investigating an environment that also has relatively high concentrations of C_2+_ gases, the potential and importance of both GoM-Arc1- and *Syntrophoarchaeaum*- like archaea degrading thermogenic hydrocarbons in cold seep sediments is shown for the first time. This points to a more widespread significance for these microbial groups in marine carbon cycling, in different environments and at different temperatures.

For anaerobic hydrocarbon oxidation to be energetically favorable, it must be coupled to the reduction of an electron acceptor either directly or through electron transfer to a syntrophic partner. None of the canonical terminal reductases (e.g. for iron, sulfate, nitrate, nitrite reduction) reported in other studies^23^ were detected in MAGs representing alkane-oxidizing archaea. This suggests a syntrophic partner organism is necessary to enable growth on short-chain hydrocarbons. Various sulfate reducers were detected in the same depths as the ANME archaea, and are likely syntrophic partners, including members of the well-known SEEP-SRB1 lineage^12^ (this group was prevalent in both metagenomes and amplicon libraries). ANME-1 MAGs were 60 to 91% complete, so it cannot be ruled out that they harbour canonical or alternative reductases e.g. as in the recently proposed bacterium-independent anaerobic methane oxidation^10^ whereby reverse methanogenesis is coupled to reduction of elemental sulfur and polysulfide. Consistent with this model, genes encoding sulfide:quinone oxidoreductase-like proteins in ANME-1 MAGs S3_bin4 and Co_bin174 were detected in deeper layers of the sediment where sulfate was depleted (e.g. relative abundances of 0.1 to 1.4% at 100 and 150 cmbsf). Similarly, most potential alkane-degrading bacterial MAGs lack genes for respiration. Members of the *Syntrophobacterales*, such as *Smithella* and *Syntrophus*, have been shown to be able to degrade alkanes by addition to fumarate under methanogenic conditions in syntrophic association with methanogens^8, 55^. It is therefore likely that both archaea and bacteria rely on syntrophic partners to complete anaerobic hydrocarbon degradation, not only in terms of metabolite trade-offs, but also for accepting electrons^56, 57^.

Besides hydrocarbons supplied from deep thermogenic petroleum deposits, metabolic reconstructions of 376 MAGs also revealed versatile catabolic capabilities for assimilating carbohydrates, peptides and short chain lipids. This indicates that the utilization of various types of sedimentary organic carbon (e.g. recent organic matter deposited with the sediments) is important to the biogeochemistry of deep sea sediments, including at cold seeps that also receive inputs of thermogenic hydrocarbons from below^4, 58–60^. Also widespread is the capacity to conserve energy and fix carbon using electrons derived from inorganic compounds such as sulfide, thiosulfate, hydrogen, ammonia, and nitrite. With respect to electron acceptors, it is generally thought that sulfate reduction predominates in marine sediments except where higher-potential electron acceptors such as oxygen or nitrate are available^61, 62^, or at depths where sulfate is depleted. While bacteria capable of respiring sulfate, oxygen, nitrate, and nitrite were detected, these capacities were not prevalent throughout the community. Instead, many genomes encoded putative reductive dehalogenase genes for organohalide respiration. Diverse organohalide compounds can be produced through abiotic and biotic processes in the marine subsurface and are among the highest-potential oxidants in anoxic ecosystems (*E*°’ = 0.24 to 0.58 V). These compounds may serve as important respiratory electron acceptors for dehalogenating bacteria and archaea^4, 63^. As suggested by recent studies^63, 64^, the ability to dehalogenate may provide a competitive advantage that allows organisms to persist in subsurface sediments. In line with other recent observations^4^, fermentation appears to be a universal strategy in these sediments, resulting in hydrogen, formate, and acetate being major end products; accordingly, the majority of the recovered MAGs harboured the bidirectional [NiFe]-hydrogenase. Fermentation products are likely to serve as energy sources for other community members harbouring respiratory hydrogenases, formate dehydrogenases, and the Wood-Ljungdahl pathway.

Microbial communities and activities at this newly discovered NW Atlantic deep sea study site demonstrate that migrated thermogenic hydrocarbons sustain diverse microbial populations spanning different redox regimes in cold seep sediments. These include key hydrocarbon degraders, syntrophic partners, and other community members through interspecies transfer of electrons and metabolites, and degradation of necromass^20, 65^. These findings indicate that upward migrated thermogenic hydrocarbons are important carbon and energy sources that sustain diverse subseafloor microbial communities at permanently cold hydrocarbon seeps.

## Methods

### Sampling and geochemical characterization

This study provides a detailed analysis of a 3.44-meter-long piston core taken from the seabed of the Scotian slope (43.010478 N, 60.211777 W) in 2306 m water depth. Subsamples were collected immediately from the base of the core and stored in gas-tight isojars flushed with N_2_ for headspace gas analysis. Multiple depths ranging from the deepest portion to within about a metre or so of the top of the core, were subsampled for geochemical analysis. Additional intervals were preserved for separate microbiological analyses. Detailed subsampling depths can be found in **Tables 1 and 2** as well as **Supplementary Table 1**.

Hydrocarbon compositions of headspace gas samples were analysed using an Agilent 7890A RGA Gas Chromatograph equipped with Molsieve and Poraplot Q columns and a flame ionisation detector. Stable carbon and hydrogen isotopic signatures were determined by Trace GC2000 equipped with a Poraplot Q column, connected to a Thermo Finnigan Delta plus XP isotope ratio mass spectrometer (IRMS). Sediment samples were analysed for TOC and EOM using an Agilent 7890A RGA Gas Chromatograph equipped with a CP-Sil-5 CB-MS column. A Micromass ProSpec-Q instrument was used for determination of saturated and aromatic fractions. Stable carbon isotope analyses of these fractions were determined on a Eurovector EA3028 connected to a Nu Horizon IRMS. Experimental procedures on these measurements followed ‘The Norwegian Industry Guide to Organic Geochemical Analyses, Edition 4.0 (30 May 2000)’.

### Sulfate measurement

Porewater sulfate concentrations were measured in a Dionex ICS-5000 reagent-free ion chromatography system (Thermo Scientific, CA, USA) equipped with an anion-exchange column (Dionex IonPac AS22; 4 × 250 mm; Thermo Scientific), an EGC-500 K_2_CO_3_ eluent generator cartridge and a conductivity detector.

### Metabolomic analysis

Pore water metabolites were extracted from sediment samples according to previously reported methods^4^. Mass spectrometric (MS) analysis was carried out using a Thermo Scientific Q-Exactive^TM^ HF Hybrid Quadrupole-Orbitrap mass spectrometer with an electrospray ionization source coupled to ultra high-performance liquid chromatography (UHPLC). Data were acquired in negative ion mode using full scan from 50-750 *m/z* at 240,000 resolution with an automatic gain control (AGC) target of 3e^6^ and a maximum injection time of 200 ms. For MS/MS fragmentation, an isolation window of 1 *m/z* and an AGC target of 1e^6^ was used with a maximum injection time of 100 ms. Data were analyzed for specific *m*/*z* ratios using MAVEN software^66^.

### 16S rRNA gene amplicon sequencing

DNA was extracted from sediment samples using the PowerSoil DNA Isolation Kit (12888-50, QIAGEN). Amplification of the v3-4 region of the bacterial 16S rRNA genes and the v4-5 region of the archaeal 16S rRNA genes, used primer pairs SD-Bact-0341-bS17/SD-Bact-0785-aA21 and SD-Arch-0519-aS15/SD-Arch-0911-aA20, respectively^67^. Amplicon sequencing was performed on a MiSeq benchtop sequencer (Illumina Inc.) using the 2 × 300 bp MiSeq Reagent Kit v3. Reads were quality controlled and then clustered into operational taxonomic units (OTUs) of >97% sequence identity with MetaAmp^68^. Taxonomy was assigned using the SILVA database^69^ (release 132).

### Quantitative PCR

Quantitative polymerase chain reaction (qPCR) analyses were performed on the new DNA extracts using PowerSoil DNA Isolation Kit (12888-50, QIAGEN) to estimate the abundance of bacteria and archaea at different depths in the sediment core. The mass of sediment used for DNA extraction was typically 0.5 g and was always recorded. PCR reactions were set up using Bio-Rad SsoAdvanced Universal SYBR Green Supermix. Amplification of bacterial and archaeal 16S rRNA genes used domain-specific primers B27F-B357R and A806F-A958R, respectively. Triplicate PCR was performed on a Thermo Scientific PikoReal Real-Time PCR Instrument, using reaction conditions described previously^70^. Results were recorded and analysed by PikoReal software 2.2.

### Metagenome sequencing

DNA was extracted from the sediment samples using the larger format PowerMax Soil DNA Isolation Kit (12988-10, QIAGEN) according to the manufacturer’s instructions. Metagenomic library preparation and DNA sequencing using NextSeq 500 System (Illumina Inc.) were conducted at the Center for Health Genomics and Informatics in the Cumming School of Medicine, University of Calgary.

### Microbial diversity analysis

SingleM (https://github.com/wwood/singlem) was applied to raw metagenome reads from each sample^30^. Shannon diversity was calculated based on SingleM counts on 14 single copy marker genes. The vegan package was then used to calculate diversity based on the rarefied SingleM OTU table across each of the 14 marker genes. The average was taken as the Shannon index determination for each sample. To explore microbial composition of each sample, full-length 16S rRNA genes were reconstructed from metagenomic raw reads using the phyloFlash pipeline^31^ together with SILVA database^69^ (release 132).

### Assembly and binning

Raw reads were quality-controlled by (1) clipping off primers and adapters and (2) filtering out artifacts and low-quality reads using the BBDuk function of BBTools (https://sourceforge.net/projects/bbmap/). Filtered reads were co-assembled using MEGAHIT^71^ and were individually assembled using metaSPAdes^72^. For co-assembly, one additional metagenome (315 cmbsf) sequenced using the same method was also included. This depth was discarded for discussion as it might be contaminated with seawater (**Supplementary Note 1**). Short contigs (<1000 bp) were removed from assemblies. For each assembly, binning used the Binning module within metaWRAP^35^ (--maxbin2 --metabat1 --metabat2 options). Resulting bins were then consolidated into a final bin set with metaWRAP’s Bin_refinement module (-c 50 -x 10 options). All binning results were combined and dereplicated using dRep^73^ (-comp 50 -con 10 options). After dereplication, a total of 376 dereplicated MAGs were obtained.

### Calculating relative abundances and replication rates

For producing indexed and sorted BAM files, quality-controlled reads from each sample were mapped to the set of dereplicated genomes using BamM v1.7.3 ‘make’ (https://github.com/Ecogenomics/BamM). To calculate relative abundance of each MAG within a microbial community in a given sediment depth, CoverM v0.3.1 ‘genome’ (https://github.com/wwood/CoverM) was used to obtain relative abundance of each genome (parameters: --min-read-percent-identity 0.95 --min-read-aligned-percent 0.75 --trim-min 0.10 --trim-max 0.90).

Microbial replication rates were estimated with iRep^37^ for high-quality dereplicated MAGs (>=75% complete, <=175 fragments/Mbp sequence, and <=2% contamination). Replication rates were retained only if they passed the default thresholds: min cov. = 5, min wins. = 0.98, min r^2 = 0.9, GC correction min r^^2^ = 0.0. The require ordered SAM files were generated using the Bowtie2 (--reorder flag)^74^.

### Functional annotations

To compare abundances of metabolic genes at different sediment depths, all quality-controlled reads were aligned against comprehensive custom databases^61^ using DIAMOND BLASTx^75^ (cutoffs: e-value:1e-10, identity: 70%; best hits reserved).

For contigs, gene calling was performed using Prodigal (-p meta)^76^. Proteins were predicted against the KEGG database using GhostKOALA^48^ and against the Pfam and TIGRfam HMM models using MetaErg^77^. For individual MAGs, completeness of various metabolic pathways was determined using KEGG-Decoder^78^ and KEGG-Expander (https://github.com/bjtully/BioData/tree/master/KEGGDecoder). Annotations of key metabolic genes were also confirmed by phylogenetic analyses as described below.

Genes involved in anaerobic hydrocarbon degradation were screened using BLASTp (cutoffs: evalue 1e-20 + pident 30% + qcovs 70%) against local protein databases^4^. After removal of short sequences, they were further manually curated using BLASTp against NCBI-based nr protein sequences by checking top hits to relevant genes. For identification of McrA and DsrA, protein sequences were screened against local protein databases^79^ using BLASTp (cutoffs: evalue 1e-20 + pident 30% + qcovs 70%). McrA and DsrA protein sequences were cross-checked against MetaErg annotations and phylogenetic analyses, while hydrogenases were confirmed and classified using the HydDB tool^52^. The dbCAN2 web server^80^ was used for carbohydrate-active gene identification based on retaining proteins found by at least two of the tree tools (HMMER + DIAMOND + Hotpep) for further analysis.

### Taxonomy assignments of MAGs

Taxonomy assessment of MAGs was initially performed by identification of 16S rRNA genes using anvi’o^81^. Predicted sequences (only 52 in 367 MAGs) were aligned and classified using SILVA ACT^82^. Subsequently the taxonomy of each MAG was temporally assigned using GTDB-Tk^83^ (using GTDB R04-RS89). Phylogenetic trees were reconstructed based on concatenation of 43 conserved single-copy genes using RAxML^84^ settings: raxmlHPC-HYBRID -f a -n result -s input -c 25 -N 100 -p 12345 -m PROTCATLG -x 12345, as reported previously^4^. Bacterial and archaeal reference genomes were downloaded from NCBI GenBank. Finally, MAGs were identified to appropriate taxonomic levels according to the NCBI Taxonomy database taking results of all three of the above methods into account.

### Phylogenetic analysis of metabolic genes

For *mcrA*, gene sequences were aligned using the MUSCLE algorithm^85^ (-maxiters 16) and trimmed using TrimAL^86^ with parameters: -automated1. A maximum-likelihood phylogenetic tree was built using IQ-Tree^87^, parameters: -st AA -m LG+C60+F+G -bb 1000 -alrt 1000 -nt 20. For other key metabolic genes, sequences were aligned using the ClustalW algorithm included in MEGA7^88^. All alignments were manually inspected. For amino acid sequences of the group 3 [NiFe]-hydrogenase large subunit, a neighbour-joining tree was constructed using the Poisson model with gaps treated with pairwise deletion, bootstrapped with 50 replicates and midpoint-rooted. For amino acid sequences of other genes, maximum-likelihood trees of were constructed using the JTT matrix-based model with all sites, bootstrapped with 50 replicates and midpoint-rooted.

### Data availability

DNA sequences have been deposited in NCBI BioProject databases with accession number PRJNA598277 (https://www.ncbi.nlm.nih.gov/bioproject/). The authors declare that all other data supporting the findings of this study are available within the article and its supplementary information files, or from the corresponding authors upon request.

## Acknowledgements

The work was supported by Genome Canada Genomics Applications Partnership Program (GAPP) and Canada Foundation for Innovation (CFI-JELF 33752) awards to C.R.J.H., who is supported by a Campus Alberta Innovates Program Chair. X.D. is supported by National Natural Science Foundation of China (Grant No. 41906076) and the Fundamental Research Funds for the Central Universities (Grant No. 19lgpy90). Metabolomics data were obtained at the Calgary Metabolomics Research Facility (CMRF), which is supported by the International Microbiome Centre and the Canada Foundation for Innovation (CFI-JELF 34986) awards to I.A.L., who is supported by an Alberta Innovates Translational Health Chair. C.G. is supported by an ARC DECRA Fellowship (DE170100310) and an ARC Discovery Project (DP180101762). We thank Weiling Pi, Chuwen Zhang, Zexin Li and Haoyu Lan for help with figure preparation, Ben Woodcroft for help with CoverM software, and Rhonda Clark for research support.

## Author contributions

C.R.J.H. obtained the funding for this project. X.D. and C.R.J.H. designed the study. X.D. processed metagenome data. J.E.R., S.M., R.A.G. and I.A.L. performed metabolomics analyses and data interpretation. D.C.C. was the chief scientist aboard the CCGS *Hudson* and was responsible for sediment sampling. M.F., J.W. and A.M. designed and performed petroleum geochemical analyses. C.G. performed phylogenetic analysis of key metabolic genes. C.L. conducted amplicon sequencing and analyses. A. C. performed microbial diversity analyses. O.A. performed porewater sulfate measurements. S.W. and D.M. conducted qPCR analyses. X.D., C.G. and C.R.J.H. drafted the manuscript. All authors reviewed the results and participated in the writing of the manuscript.

