## Supplementary materials for "Thermogenic hydrocarbons fuel a redox stratified subseafloor microbiome in deep sea cold seep sediments"

**List of contents**

**Supplementary Note 1**

**Supplementary Note 2**

**Supplementary Table 1** Extractable organic matter compositions of four sediment subsamples.

**Supplementary Table 2** Genome information for 376 MAGs, including (1) genome completeness and contamination estimates (determined by CheckM), (2) *in situ* replication rate measurements determined by iRep, (3) the relative abundance (%) and genome coverage (×) determined by mapping each MAG against quality-filtered metagenome reads, (4) taxonomic assignment of MAGs based on 16S rRNA gene sequences where possible, (5) taxonomic assignment of MAGs based on GTDB-tK classification, and (6) taxonomic assignment of MAGs based on NCBI taxonomy determined by a phylogenetic analysis of 43 marker genes.

**Supplementary Table 3** Statistics of metagenome assemblies. Data were generated using QUAST (default parameters). All statistics are based on contigs of size >= 500 bp, unless otherwise noted (e.g., "# contigs (>= 0 bp)" and "Total length (>= 0 bp)" include all contigs).

**Supplementary Table 4** Summary for occurrence of various functions of interest encoded within each MAG. Pathways are scaled from 0 to 1, where 1 represents that the genome contains all genes for the function of interest. Detailed gene list for each pathway can be found at: https://github.com/bjtully/BioData/blob/master/KEGGDecoder.

**Supplementary Table 5** Archaeal (a) and bacterial (b) OTU tables based on 16S rRNA gene amplicon sequencing.

**Supplementary Table 6** Details for identification of genes in genomes containing *mcrA* genes based on presence/absence matrix. Also include detailed genome statistics on these genomes.

**Supplementary Table 7** Details for identification of genes in genomes containing *assA* genes based on presence/absence matrix. Also include detailed genome statistics on these genomes.

**Supplementary Table 8** Details for identification of genes in genomes containing *bcr* genes based on presence/absence matrix. Also include detailed genome statistics on these genomes. Only MAGs contain at least two further genes in this pathway are discussed in the main text.

**Supplementary Table 9** Summary of carbohydrate-active enzymes (CAZymes) detected in 376 MAGs. Total numbers for carbohydrate esterases (CE), glycoside hydrolases GH) and polysaccharide lyases (PL), as well as corresponding raw outputs from the dbCAN2 webserver, are shown.

**Supplementary Table 10** Total number of peptidases detected in each MAG. Total number of peptidases identified using the Pfam HMM models related to MEROPS peptidase database.

**Supplementary Table 11** Details for total number of genes related to the beta oxidation pathway in each genome using the KAAS webserver.

**Supplementary Table 12** Details for (1) identification of genes encoding for different types of hydrogenases and (2) identified genes encoding reductive dehalogenases in each MAG. Hydrogenases were classified using the hydrogenase classifier HydDB while the *rdhA* genes were detected using TIGR02486.

**Supplementary Figure 1** Map of the studied sampling location. Adapted from: https://doi.org/10.4095/314695.

**Supplementary Figure 2** GC-FID chromatograms of extractable organic matter showing unresolved complex mixture (UCM) humps. Samples from four different depths were analyzed, (a) 53-60 cmbsf, (b) 208-213 cmbsf, (c) 227-232 cmbsf, and (d) 310-315 cmbsf. y-axis: detector response; x-axis: retention time (minutes). Additional parameters from the EOM for these samples are provided in **Supplementary Table 1**.

**Supplementary Figure 3** Pore water concentrations of sulfate and proposed biogeochemical zonation within the core showing a mixing zone, sulfate reduction zone, sulfate methane transition zone, and methanogenic zone.

**Supplementary Figure 4** Maximum-likelihood tree of amino acid sequences of catalytic subunits of canonical fumarate-adding enzymes, which are markers for anaerobic hydrocarbon degradation. The tree shows sequences from cold seep metagenome-assembled genomes (blue) alongside representative reference sequences (black). Different clades correspond to alkylsuccinate synthases (AssA) as well as benzylsuccinate synthases (BssA), naphthylmethylsuccinate synthases (NmsA), and hydroxybenzylsuccinate synthases (HbsA). The tree was constructed using the JTT matrix-based model, used all sites, and was bootstrapped with 50 replicates and midpoint-rooted.

**Supplementary Figure 5** Distribution of major KEGG categories at different sediment depths. Annotations were performed on contigs assembled from the quality-controlled reads of each depth. X-axis indicates the different sediment depths, and y-axis indicates major functional categories. For each category, values were normalized by their standard score (z-score).

**Supplementary Figure 6** Neighbour-joining tree of amino acid sequences of the group 3 [NiFe]-hydrogenase large subunit, a marker for bidirectional hydrogen metabolism during various processes. The tree shows sequences from cold seep metagenome-assembled genomes (blue) alongside representative reference sequences (black). The subgroup of each reference sequence is denoted according to the HydDB classification scheme. The tree was constructed using the Poisson model with gaps treated with pairwise deletion, was bootstrapped with 50 replicates, and was midpoint-rooted. To enable neighbour-joining, 54 incomplete sequences of the original 298 hits were excluded from this analysis.

**Supplementary Figure 7** Maximum-likelihood tree of amino acid sequences of flavocytochrome *c* sulfide dehydrogenase (FCC), a marker for sulfide oxidation. The tree shows sequences from cold seep metagenome-assembled genomes (blue) alongside representative reference sequences (black). The tree was constructed using the JTT matrix-based model, used all sites, and was bootstrapped with 50 replicates and midpoint-rooted.

**Supplementary Figure 8** Maximum-likelihood tree of amino acid sequences of the sulfide-quinone oxidoreductase (Sqr), a marker for sulfide oxidation. The tree shows sequences from cold seep metagenome-assembled genomes (blue) alongside representative reference sequences (black). The subgroup of each reference sequence is denoted. The tree was constructed using the JTT matrix-based model, used all sites, and was bootstrapped with 50 replicates and midpoint-rooted.

**Supplementary Figure 9** Maximum-likelihood tree of amino acid sequences of dissimilatory sulfite reductase A subunit (DsrA). The tree shows sequences from cold seep metagenome-assembled genomes (blue) alongside representative reference sequences (black). This enzyme is a marker for dissimilatory sulfite reduction (top and bottom major clades; *Deltaproteobacteria*, Zixibacteria, *Dehalococcoidia* bins) and sulfide oxidation (middle clade; *Gammaproteobacteria* and *Alphaproteobacteria* bins). The tree was constructed using the JTT matrix-based model, used all sites, and was bootstrapped with 50 replicates and midpoint-rooted.

**Supplementary Figure 10** Maximum-likelihood tree of amino acid sequences of the thiosulfohydrolase (SoxB), a marker for thiosulfate oxidation. The tree shows sequences from cold seep metagenome-assembled genomes (blue) alongside representative reference sequences (black). The subgroup of each reference sequence is denoted. The tree was constructed using the JTT matrix-based model, used all sites, and was bootstrapped with 50 replicates and midpoint-rooted.

**Supplementary Figure 11** Maximum-likelihood tree of amino acid sequences of ammonia monooxygenase A subunit (AmoA), a marker for ammonia oxidation during aerobic nitrification. The tree shows sequences from cold seep metagenome-assembled genomes (blue) alongside representative reference sequences (black). The tree was constructed using the JTT matrix-based model, used all sites, and was bootstrapped with 50 replicates and midpoint-rooted.

**Supplementary Figure 12** Maximum-likelihood tree of amino acid sequences of the nitrite oxidoreductase A subunit (NxrA), a marker for nitrite oxidation during aerobic nitrification. The tree shows sequences from cold seep metagenome-assembled genomes (blue) alongside representative reference sequences (black). The subgroup of each reference sequence is denoted. The tree was constructed using the JTT matrix-based model, used all sites, and was bootstrapped with 50 replicates and midpoint-rooted.

**Supplementary Figure 13** Maximum-likelihood tree of amino acid sequences of carbon monoxide dehydrogenase large subunit (CoxL), a marker for aerobic carbon monoxide oxidation. The tree shows sequences from cold seep metagenome-assembled genomes (blue) alongside representative reference sequences (black). The tree was constructed using the JTT matrix-based model, used all sites, and was bootstrapped with 50 replicates and midpoint-rooted.

**Supplementary Figure 14** Maximum-likelihood tree of amino acid sequences of group 1 and 2 [NiFe]-hydrogenase large subunits, a marker for hydrogen oxidation during respiratory processes. The tree shows sequences from cold seep metagenome-assembled genomes (blue) alongside representative reference sequences (black). The subgroup of each reference sequence is denoted according to the HydDB classification scheme. The tree was constructed using the JTT matrix-based model, used all sites, and was bootstrapped with 50 replicates and midpoint-rooted.

**Supplementary Figure 15** Maximum-likelihood tree of amino acid sequences of the group 4 [NiFe]-hydrogenase large subunit, a marker primarily for hydrogen production during fermentative and respiratory processes. The tree shows sequences from cold seep metagenome-assembled genomes (blue) alongside representative reference sequences (black). The subgroup of each reference sequence is denoted according to the HydDB classification scheme. The tree was constructed using the JTT matrix-based model, used all sites, and was bootstrapped with 50 replicates and midpoint-rooted.

**Supplementary Figure 16** Maximum-likelihood tree of amino acid sequences of the reductive dehalogenase A subunit (RdhA), a marker for reductive dehalogenation. The tree shows sequences from cold seep metagenome-assembled genomes (blue) alongside representative reference sequences (black). The subgroup of each reference sequence is denoted. The tree was constructed using the JTT matrix-based model, used all sites, and was bootstrapped with 50 replicates and midpoint-rooted.

**Supplementary Data 1** Newick tree file for the maximum likelihood phylogenetic tree of 376 MAGs based on 43 concatenated protein-coding genes **in Figure 1**. Reference genomes for relatives were accessed from NCBI GenBank. The tree was built using RAxML with the PROTGAMMALG model.

**Supplementary Data 2** Sequences for amino acids that used to construct *mcrA* trees in **Figure 3**.

**Supplementary References**

**Supplementary Note 1**

Pore-water sulfate concentrations were measured at different sediment depths **(Table 2 and** **Supplementary Figure 3)**. They showed maxima at around ~28 mM in the uppermost sediments, with concentrations that decreased with depth **(Supplementary Figure 3)**, consistent with sulfate reduction coupled to the biodegradation of hydrocarbons. By 100 cmbsf, sulfate was entirely depleted. These observations together with geochemical data for hydrocarbon gases **(Tables 1 and 2)** and considerations of similar geochemical settings for other cold seeps^1, 2^, are consistent with classifying the different depths as: (1) a mixing zone at the interface between sediments and seawater; (2) the underlying sulfate reduction zone where high amounts of sulfate are observed; (3) a shallower sulfate-methane transition zone from 60 to 100 cmbsf where normally high activities of sulfate-dependent methane oxidation can be observed; (4) the methanogenic zone where sulfate is depleted **(Table 2 and Supplementary Figure 3)**. Increases in sulfate in deeper samples (>250 cmbsf) coincided with increases in other ions found in seawater and were believed to be due to seawater intrusion during piston core processing (data not shown).

**Supplementary Note 2**

To make a functional characterization of the microbial community associated with the metagenomes, we annotated the derived individual metagenome assemblies (**Supplementary Table 3**) against the KEGG database (**Supplementary Figure 5**). Clustering of level 1 KEGG functional categories revealed a marked discontinuity between surface samples and the other subsurface samples, in agreement with the taxonomic profiling (**Figure 1a**) and the redox zonation (**Supplementary Figure 3**). After comparing the number of proteins assigned to each subsystem, we found the most abundant genes to be those involved in genetic information processing, signaling and cellular processes, and carbohydrate metabolism (**Supplementary Figure 5**). Results suggest that most of the microorganisms in this environment are geared toward self-maintenance and environmental adaptation, consistent with a recent study showing that microorganisms inhibiting in subseafloor adapted to challenging environmental conditions^3^.

**Supplementary Table 1** Extractable organic matter compositions of four sediment subsamples.

| Depth (cmbsf) | | 53-60 | 208-213 | 227-232 | 310-315 |
| --- | --- | --- | --- | --- | --- |
| TOC (%) | | 0.25 | 0.35 | 0.36 | 0.56 |
| EOM (mg/kg rock) | | 104 | 361 | 177 | 168 |
| wt% of EOM | SAT | 47.5 | 52.1 | 35.0 | 24.6 |
|  | ARO | 10.0 | 11.8 | 13.9 | 12.5 |
|  | POL | 42.5 | 26.5 | 17.9 | 36.4 |
|  | ASP | 0.0 | 9.6 | 33.2 | 26.4 |
| δ¹³C-Sat (‰) | | ND | -31.5 | ND | ND |
| δ¹³C-Aro (‰) | | ND | -29.6 | ND | ND |

Abbreviations: TOC, total organic carbon; EOM, extractable organic matter; SAT, saturated hydrocarbons; ARO, aromatic hydrocarbons; POL, polars; ASP, asphaltenes; wt%, weight percentage; ND, not determined.

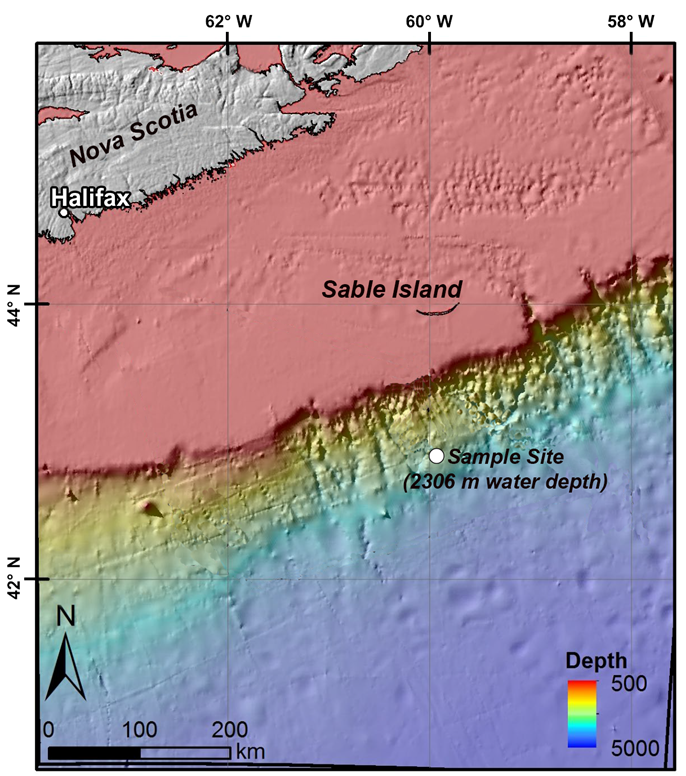

**Supplementary Figure 1** Map of the studied sampling location. Adapted from: https://doi.org/10.4095/314695.

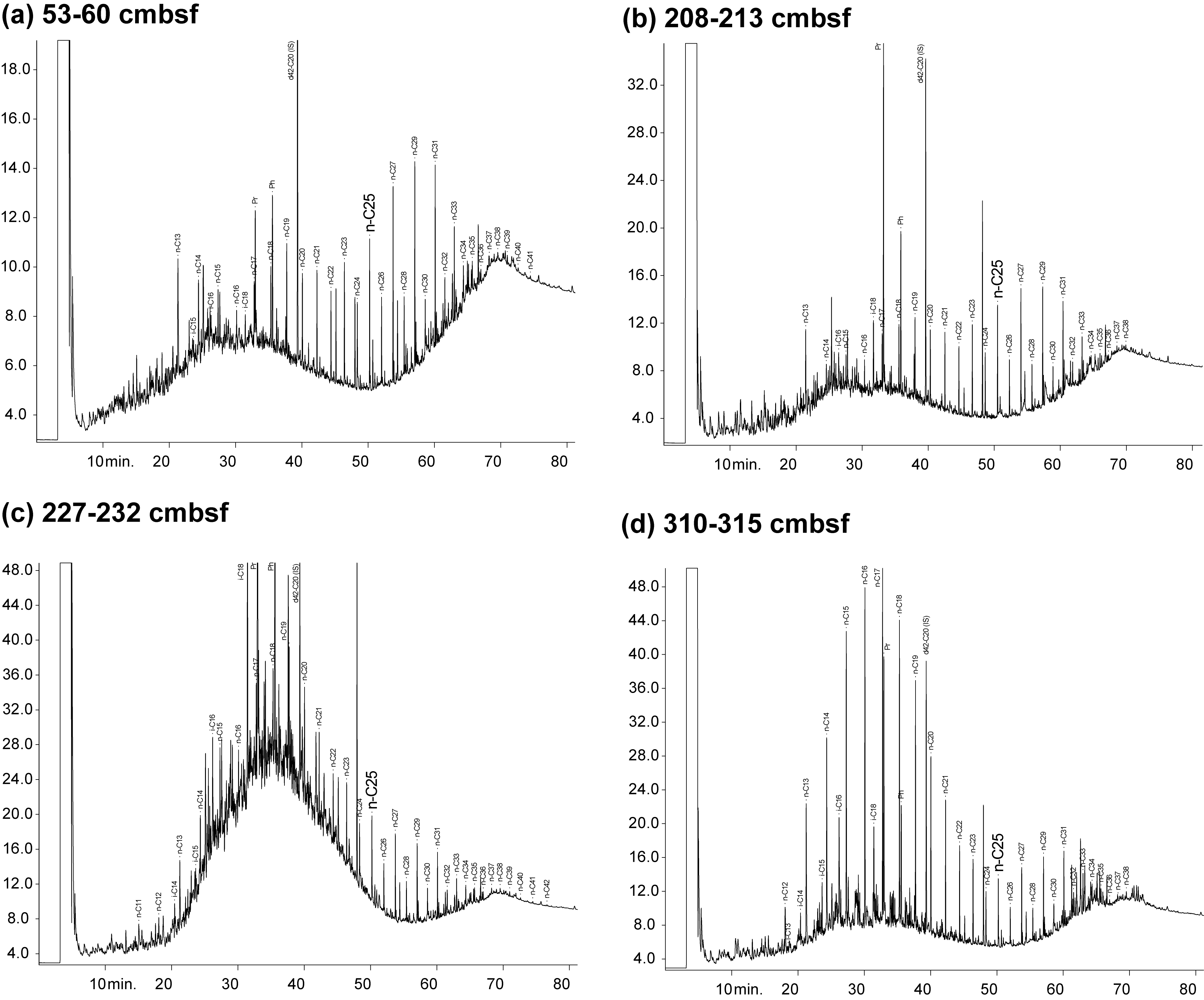

**Supplementary Figure 2** GC-FID chromatograms of extractable organic matter showing unresolved complex mixture (UCM) humps. Samples from four different depths were analyzed, (a) 53-60 cmbsf, (b) 208-213 cmbsf, (c) 227-232 cmbsf, and (d) 310-315 cmbsf. y-axis: detector response; x-axis: retention time (minutes). Additional parameters from the EOM for these samples are provided in **Supplementary Table 1**.

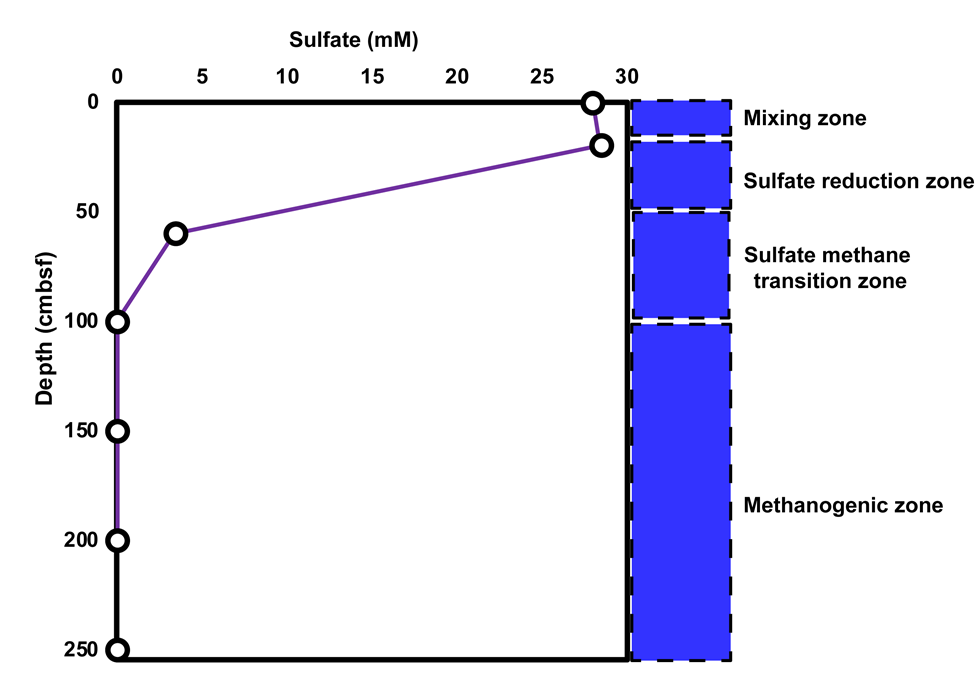

**Supplementary Figure 3** Pore water concentrations of sulfate and proposed biogeochemical zonation within the core showing a mixing zone, sulfate reduction zone, sulfate methane transition zone, and methanogenic zone.

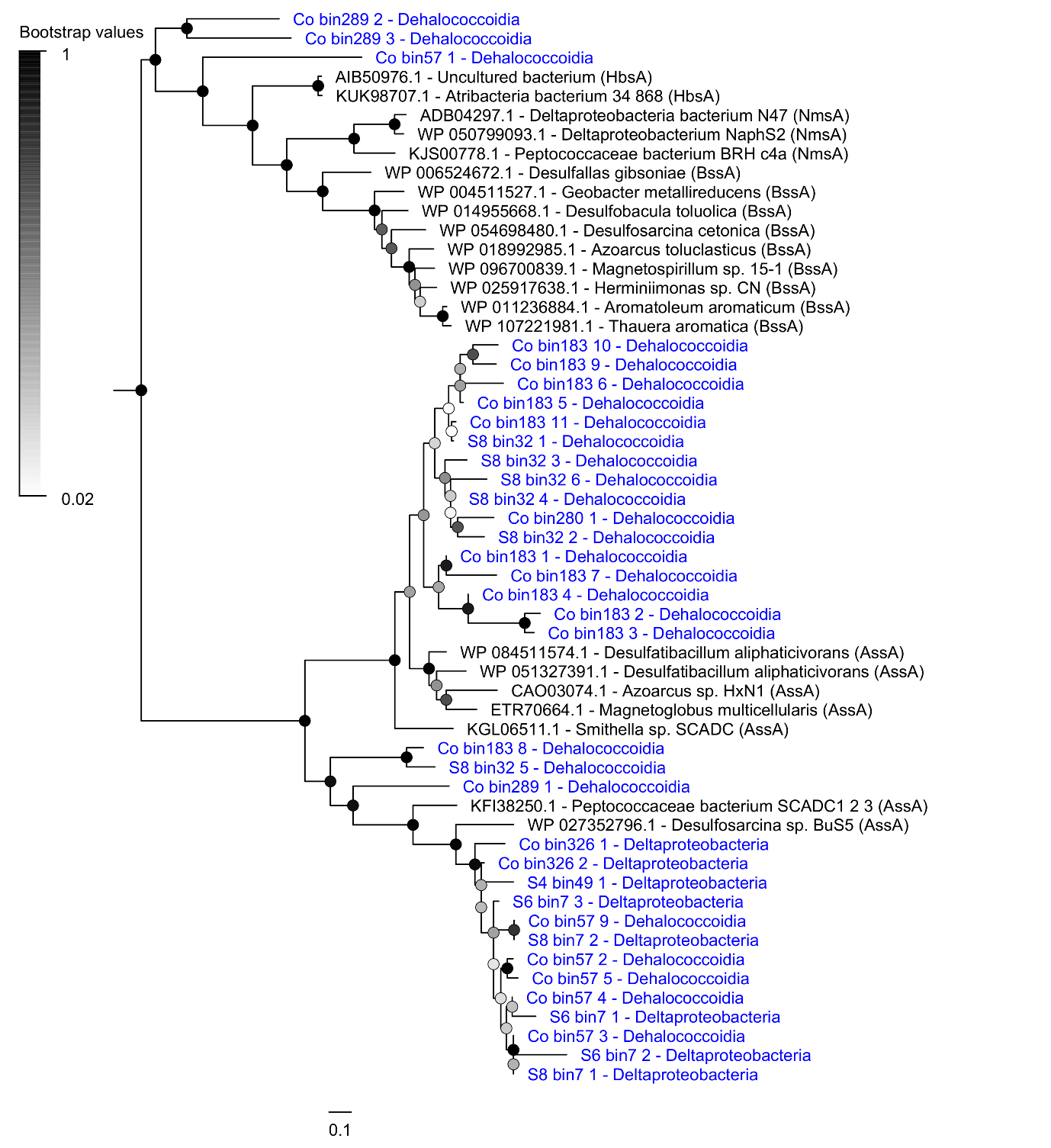

**Supplementary Figure 4** Maximum-likelihood tree of amino acid sequences of catalytic subunits of canonical fumarate-adding enzymes, which are markers for anaerobic hydrocarbon degradation. The tree shows sequences from cold seep metagenome-assembled genomes (blue) alongside representative reference sequences (black). Different clades correspond to alkylsuccinate synthases (AssA) as well as benzylsuccinate synthases (BssA), naphthylmethylsuccinate synthases (NmsA), and hydroxybenzylsuccinate synthases (HbsA). The tree was constructed using the JTT matrix-based model, used all sites, and was bootstrapped with 50 replicates and midpoint-rooted.

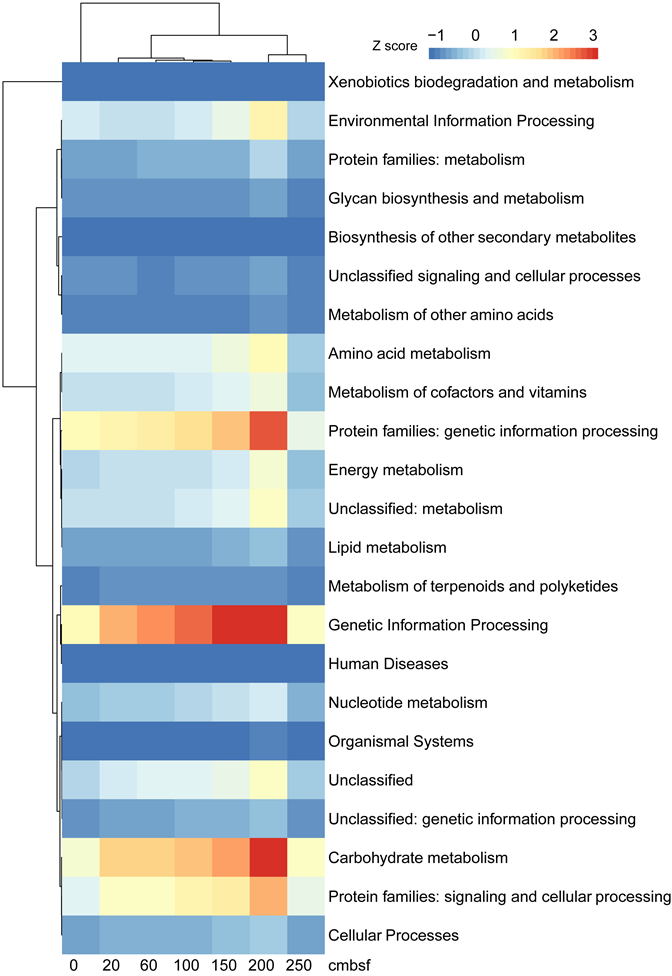

**Supplementary Figure 5** Distribution of major KEGG categories at different sediment depths. Annotations were performed on contigs assembled from the quality-controlled reads of each depth. X-axis indicates the different sediment depths, and y-axis indicates major functional categories. For each category, values were normalized by their standard score (z-score).

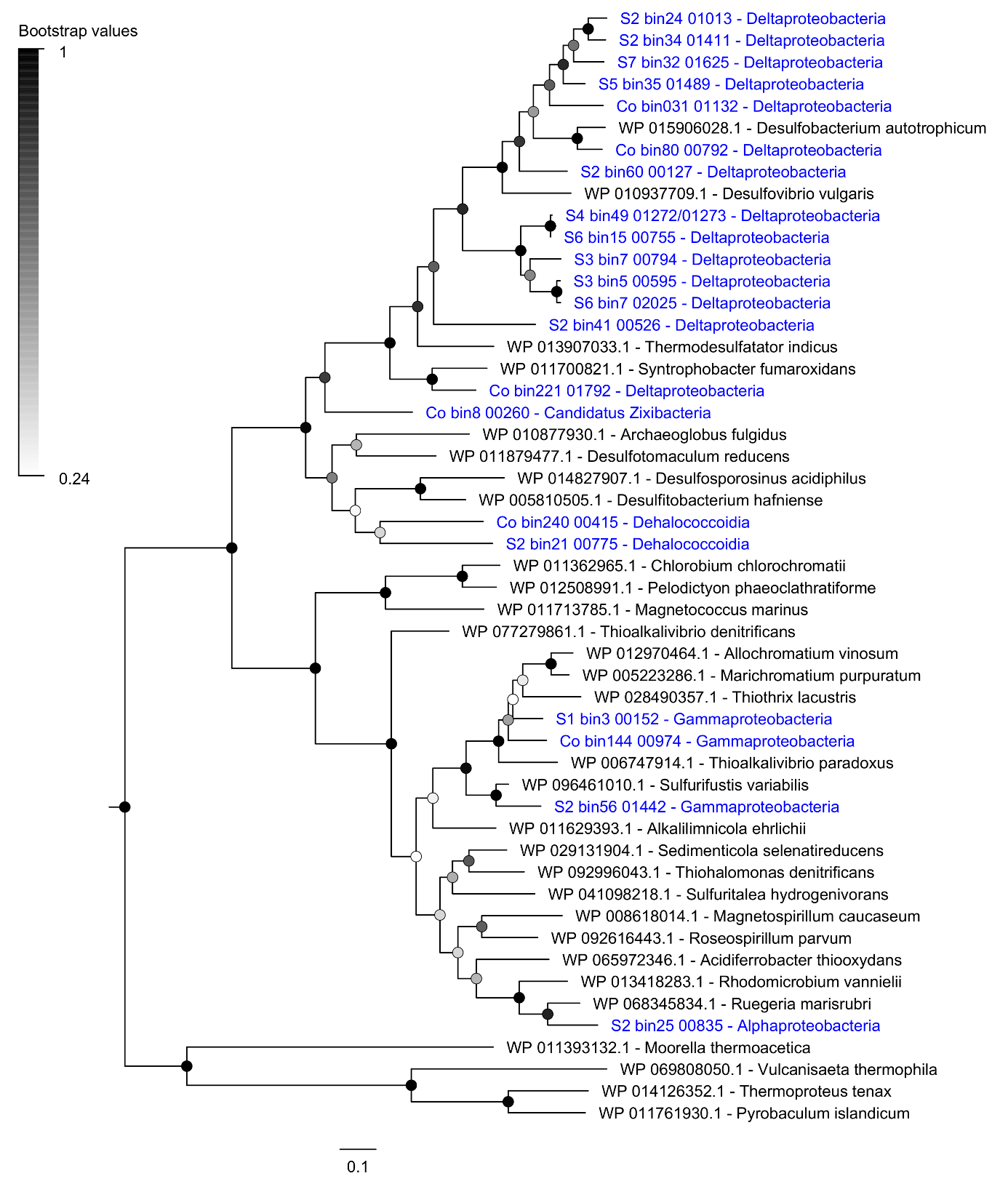

**Supplementary Figure 9** Maximum-likelihood tree of amino acid sequences of dissimilatory sulfite reductase A subunit (DsrA). The tree shows sequences from cold seep metagenome-assembled genomes (blue) alongside representative reference sequences (black). This enzyme is a marker for dissimilatory sulfite reduction (top and bottom major clades; *Deltaproteobacteria*, Zixibacteria, *Dehalococcoidia* bins) and sulfide oxidation (middle clade; *Gammaproteobacteria* and Alphaproteobacteria bins). The tree was constructed using the JTT matrix-based model, used all sites, and was bootstrapped with 50 replicates and midpoint-rooted.

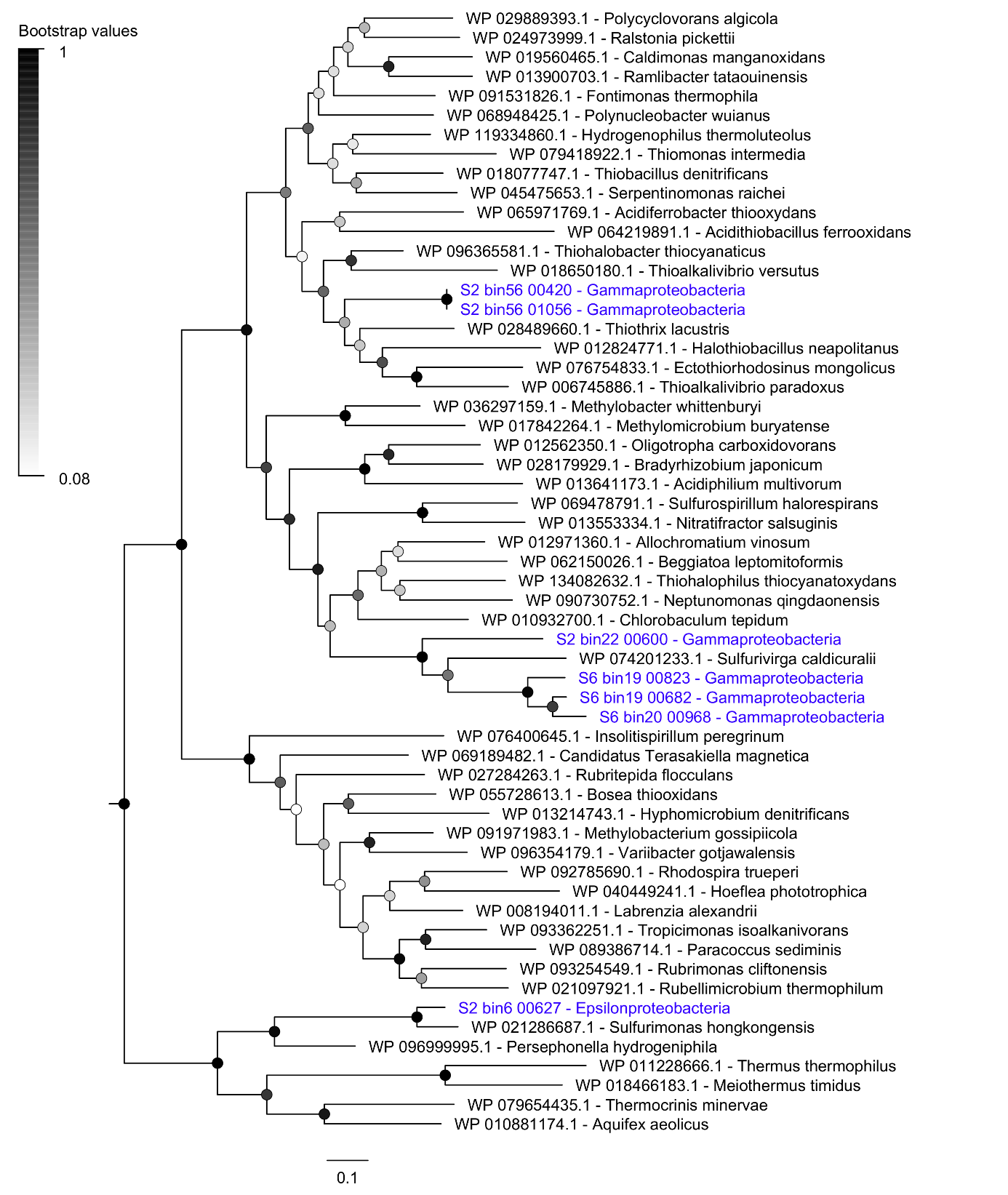
**Supplementary Figure 10** Maximum-likelihood tree of amino acid sequences of the thiosulfohydrolase (SoxB), a marker for thiosulfate oxidation. The tree shows sequences from cold seep metagenome-assembled genomes (blue) alongside representative reference sequences (black). The subgroup of each reference sequence is denoted. The tree was constructed using the JTT matrix-based model, used all sites, and was bootstrapped with 50 replicates and midpoint-rooted.

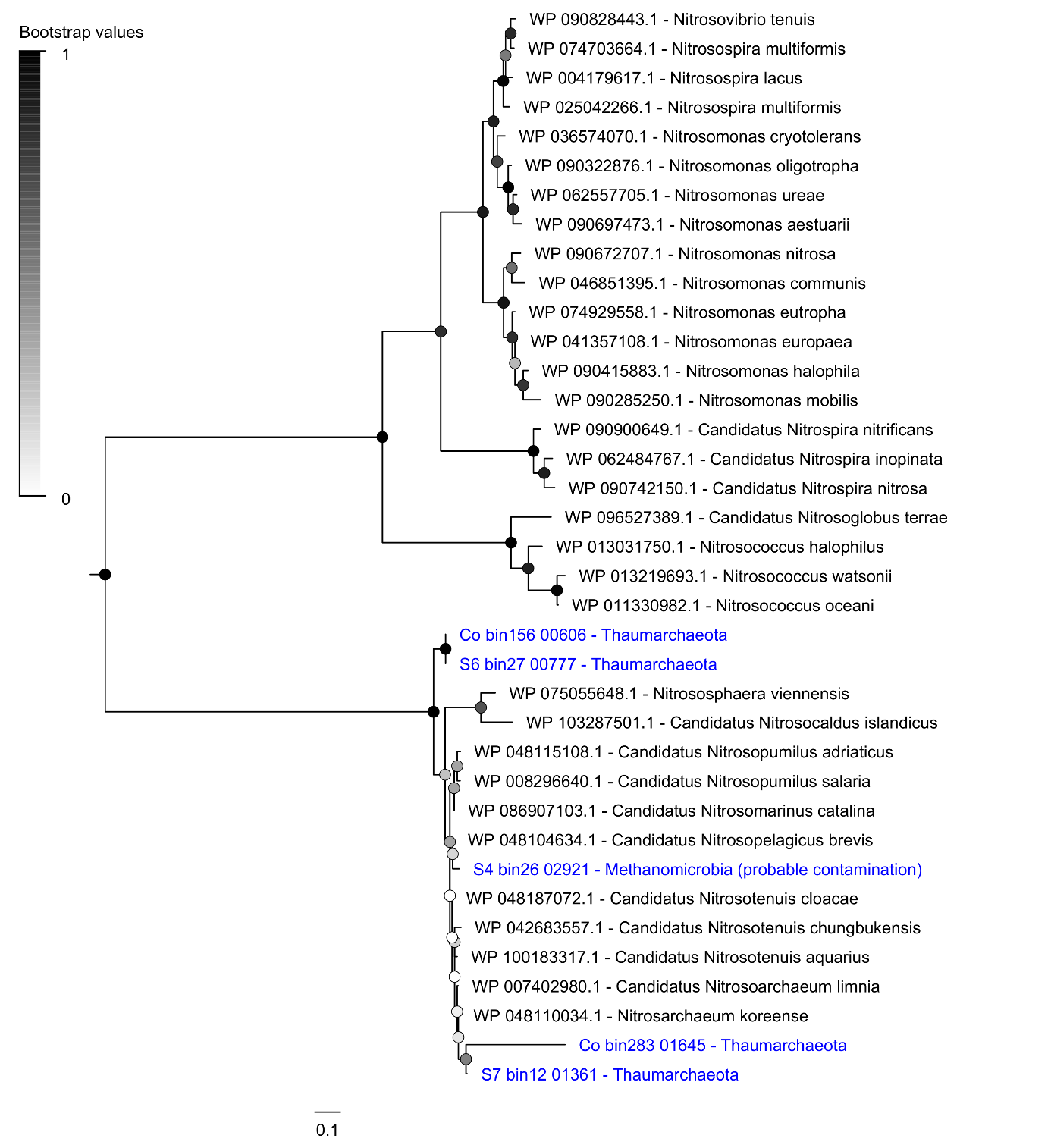
**Supplementary Figure 11** Maximum-likelihood tree of amino acid sequences of ammonia monooxygenase A subunit (AmoA), a marker for ammonia oxidation during aerobic nitrification. The tree shows sequences from cold seep metagenome-assembled genomes (blue) alongside representative reference sequences (black). The tree was constructed using the JTT matrix-based model, used all sites, and was bootstrapped with 50 replicates and midpoint-rooted.

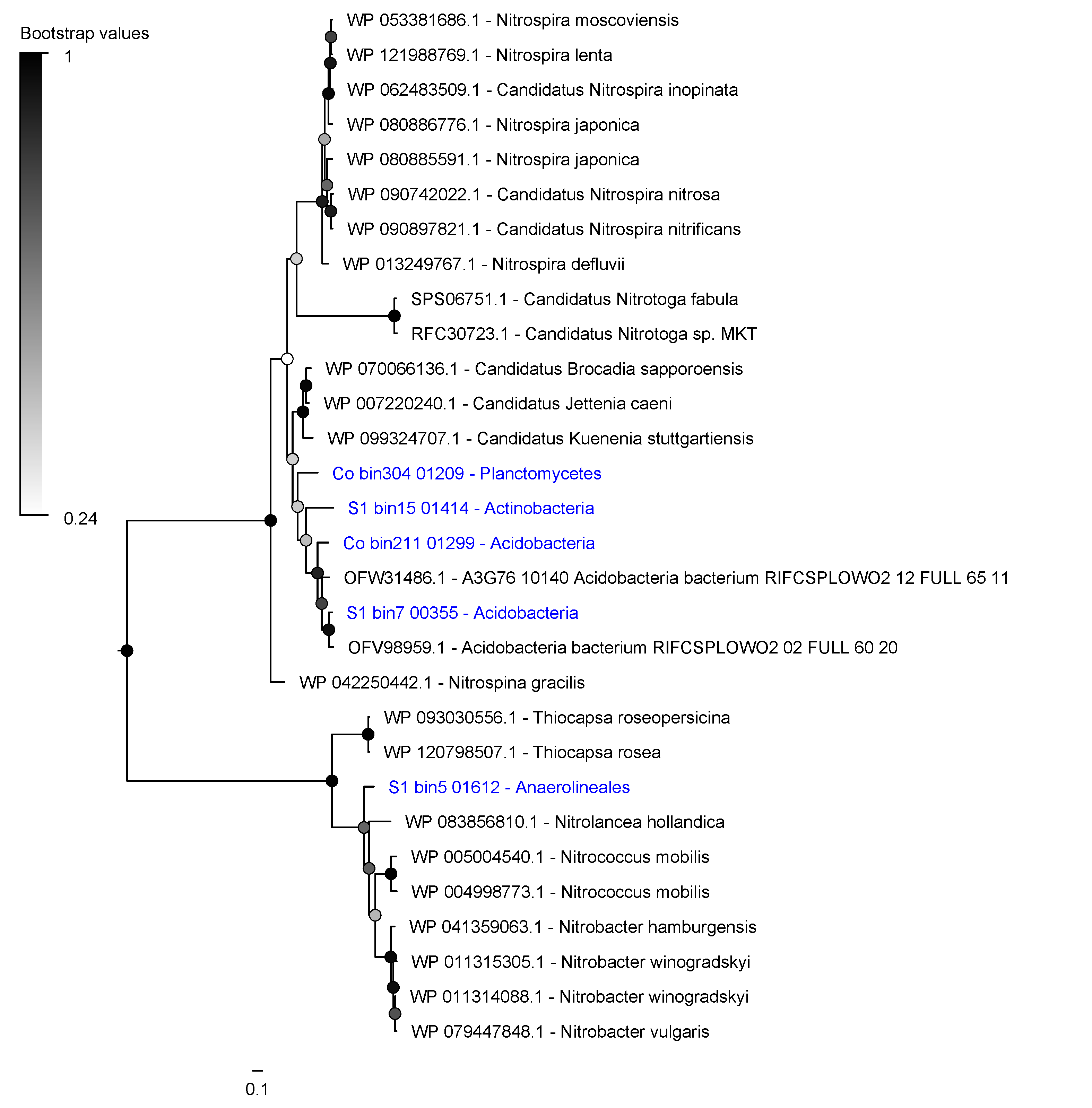
**Supplementary Figure 12** Maximum-likelihood tree of amino acid sequences of the nitrite oxidoreductase A subunit (NxrA), a marker for nitrite oxidation during aerobic nitrification. The tree shows sequences from cold seep metagenome-assembled genomes (blue) alongside representative reference sequences (black). The subgroup of each reference sequence is denoted. The tree was constructed using the JTT matrix-based model, used all sites, and was bootstrapped with 50 replicates and midpoint-rooted.

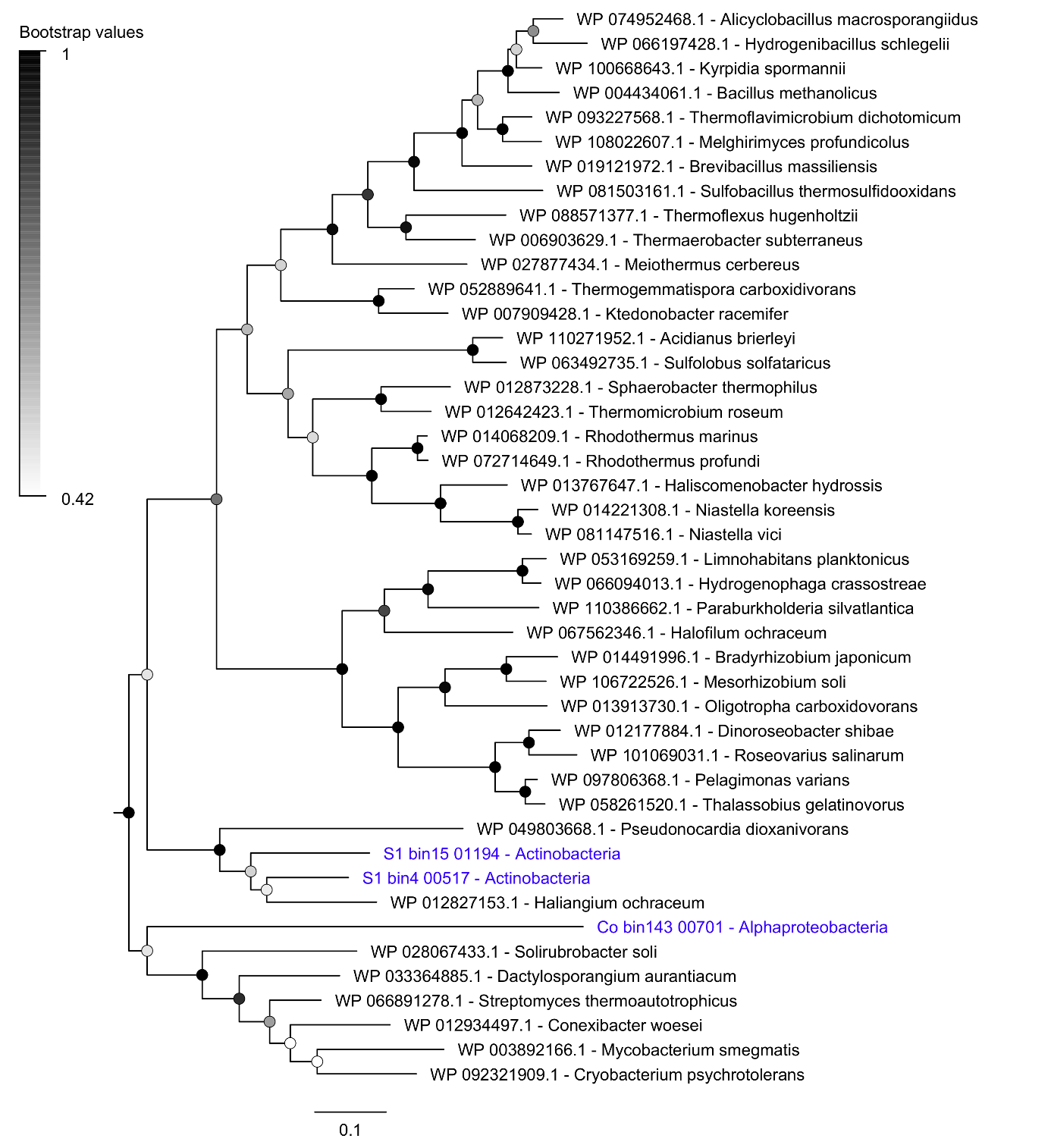
**Supplementary Figure 13** Maximum-likelihood tree of amino acid sequences of carbon monoxide dehydrogenase large subunit (CoxL), a marker for aerobic carbon monoxide oxidation. The tree shows sequences from cold seep metagenome-assembled genomes (blue) alongside representative reference sequences (black). The tree was constructed using the JTT matrix-based model, used all sites, and was bootstrapped with 50 replicates and midpoint-rooted.
