## Supplementary figures and images for "Thermogenic hydrocarbons fuel a redox stratified subseafloor microbiome in deep sea cold seep sediments"

### Supplementary Figure 6

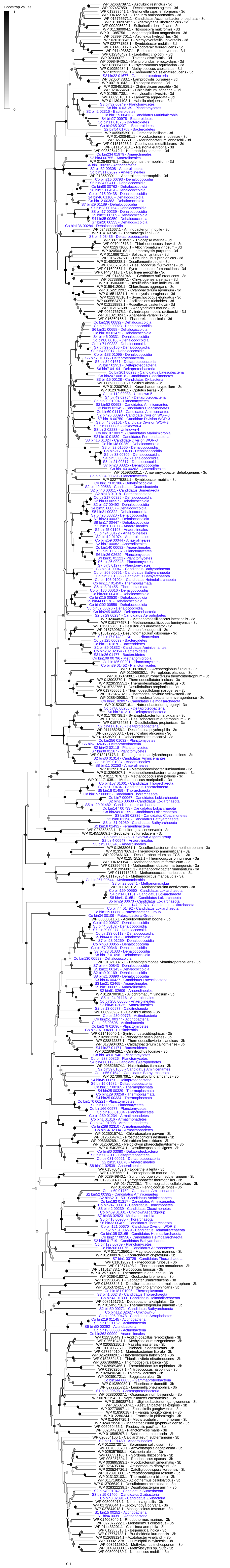

### Supplementary Figure 7

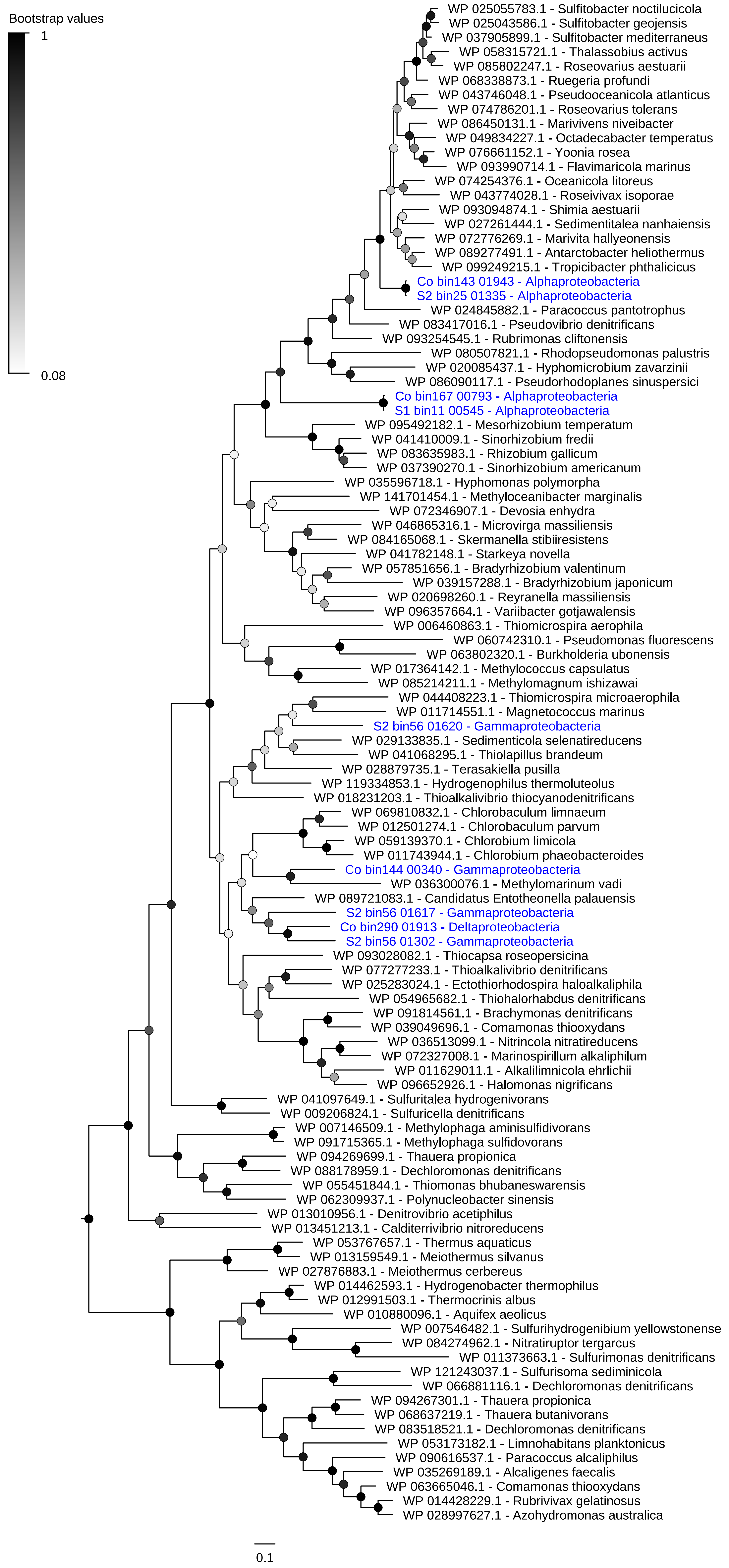

### Supplementary Figure 8

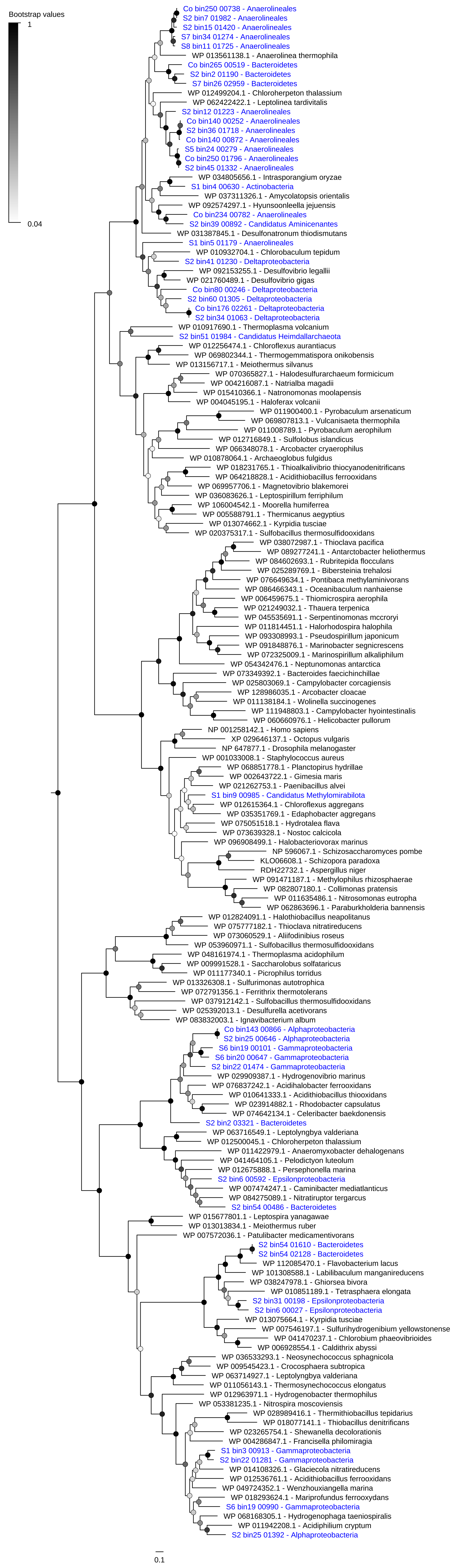

### Supplementary Figure 14

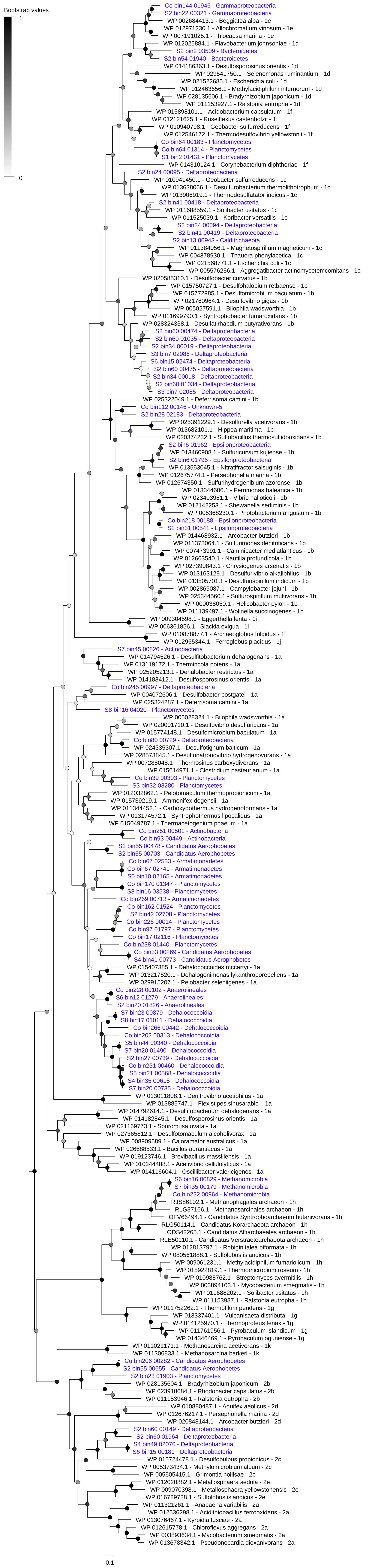

### Supplementary Figure 15

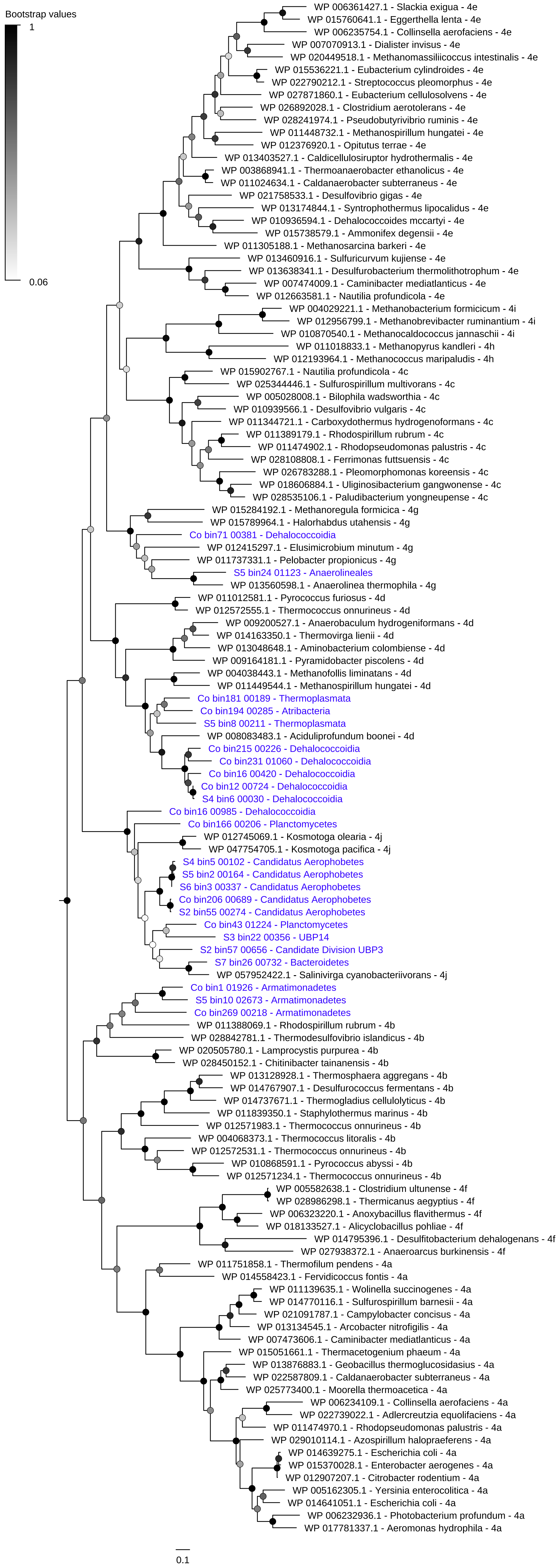

### Supplementary Figure 16

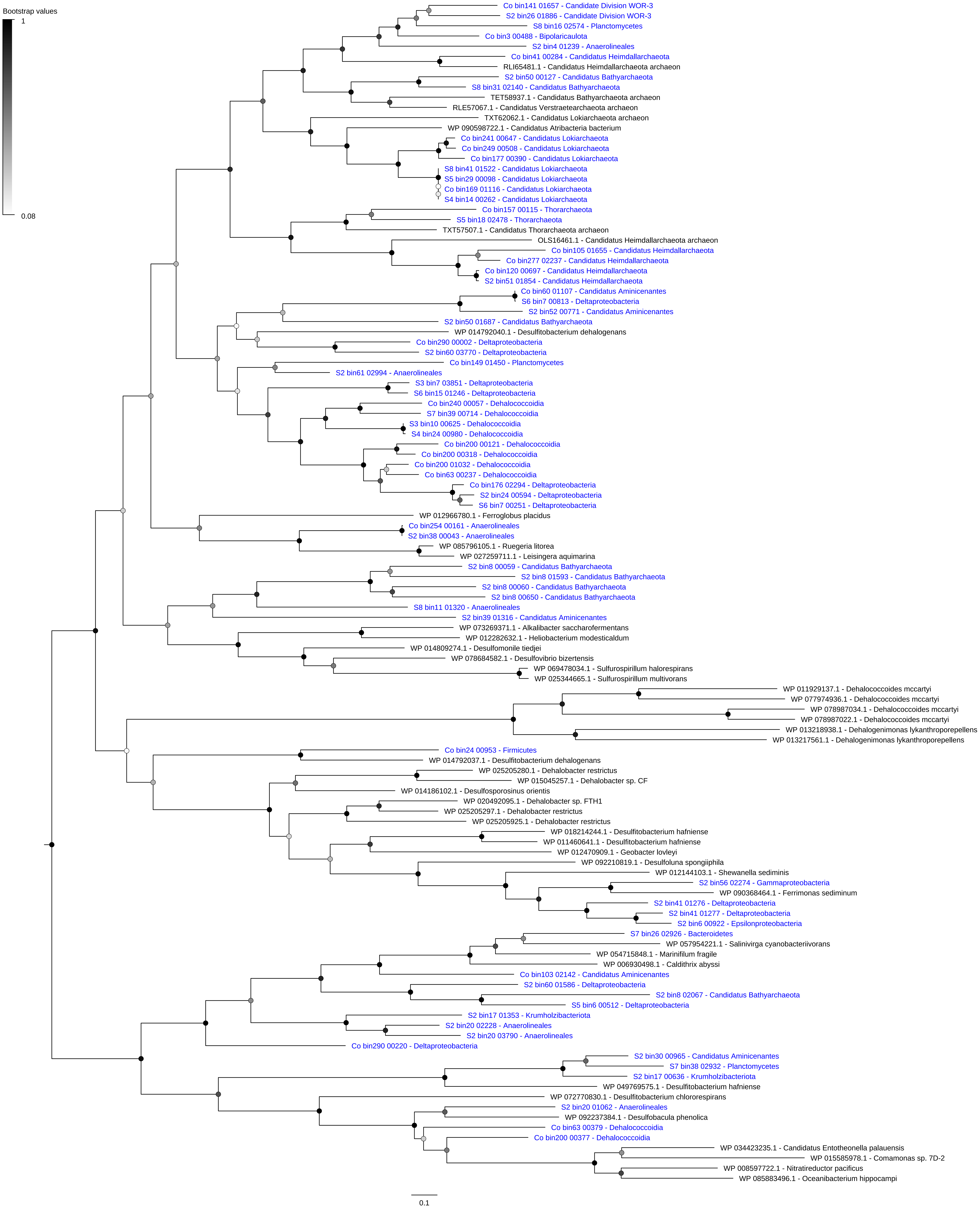
